# *O*-mannose glycosylations influence E-cadherin functional interactions

**DOI:** 10.1101/2025.08.05.668670

**Authors:** Shaoshuai Xie, Katarina Madunić, Omar G. Rosas Bringas, Weihua Tian, Sergey Y. Vakhrushev, Hjalmar Permentier, Peter Horvatovich, Adnan Halim, John LaCava

**Affiliations:** European Research Institute for the Biology of Ageing, University Medical Center Groningen, 9713 AV Groningen, NL; National Glycoengineering Research Center, Shandong University, 266237 Qingdao, CN; Department of Cellular and Molecular Medicine, Faculty of Health Sciences, Copenhagen Center for Glycomics, University of Copenhagen, DK-2200 Copenhagen, DK; Laboratory of Cellular and Structural Biology, The Rockefeller University, NY 10065, US; Department of Biotechnology and Biomedicine, Section for Medical Biotechnology, Technical University of Denmark, DK-2800 Kgs. Lyngby, DK; Department of Analytical Biochemistry, Groningen Research Institute of Pharmacy, University of Groningen, 9713 GZ Groningen, NL

**Keywords:** E-cadherin, *O* -Mannose glycosylation, TMTC enzymes, affinity proteomics

## Abstract

Cadherins are plasma membrane proteins that play critical roles in maintaining cell-cell adhesion and modulating cell signaling during development. Their functions are mediated by extracellular cadherin (EC) domains, which facilitate adhesive interactions and enable the formation of *cis-* and *trans* assemblies at adherens junctions and desmosomes. EC domains adopt a characteristic immunoglobulin-like fold composed of seven *β*-strands (A-G) and are modified by *N* -linked and *O* -linked glycosylations, including *O* -linked mannose monosaccharides (*O* -Man) on conserved serine and threonine residues of B- and G-strands. *O* -Man glycosylations on EC domains are catalyzed by the TMTC1-4 enzymes, with different TMTC enzymes modifying B- or G-strands. Given the site-specific deposition of *O* -Man glycans by dedicated enzymes and the central role of EC domains in cadherins’ functions, we hypothesized that these PTMs may fine-tune cellular adhesion and otherwise contribute to diverse physical interactions that involve cadherins. To test these hypotheses, we assayed for changes in protein-protein interactions formed with epithelial (E)-cadherin in model cells where *O* -Man were genetically ablated. Herein, we report *O* -Man-dependent E-cadherin (CDH1) protein interactions, revealed by affinity proteomics, and we orthogonally validate an altered association between CDH1 and CDH3 (P-cadherin). We show different interactomic changes associated with *O* -Man ablation on B- vs. G-strands, highlighting the importance of these PTMs in CDH1-associated interaction. These findings provide new insights into how *O* -Man regulates CDH1-dependent protein complexes.

## Introduction

The Ca^2+^-dependent cadherin superfamily of adhesion molecules comprises ∼120 members, including the classical cadherins (e.g. E-, N-, P-cadherin), atypical cadherins (e.g. cadherin-11 and cadherin-13), the clustered protocadherins (*α*-, *β*-, and *γ*-) and atypical protocadherins (e.g. PCDH15 and FAT1-4), all of which are characterized by a variable number of extracellular (EC) domains required for adhesion and function (1). Cadherins and protocadherins participate in numerous biological functions, including embryonic development (2), tissue homeostasis (3), and neuronal barcoding (4, 5); their dysregulation is associated with developmental disorders (6) and cancer (7, 8).

Within cadherins and protocadherins, EC domains form functional units that are responsible for directing Ca^2+^-dependent physical interactions within the extracellular matrix. Each EC domain is composed of seven antiparallel *β*-strands (Fig. 1A), arranged as two *β*-sheets (9–11); *β*-strands A, G, F, and C form one *β*-sheet, and *β*-strands D, E, and B form another (12). EC domains are post-translationally modified by the “<u>t</u>ransmembrane *O* -<u>m</u>annosyltransferase <u>t</u>argeting <u>c</u>adherins” enzymes (encoded by the *TMTC* genes). A multigene query of the *TMTC* genes, using the ‘Genotype Tissue Expression (GTEx) Portal,’ confirmed that they are broadly transcribed but differentially regulated across tissues (13–15).

**Figure 1.**
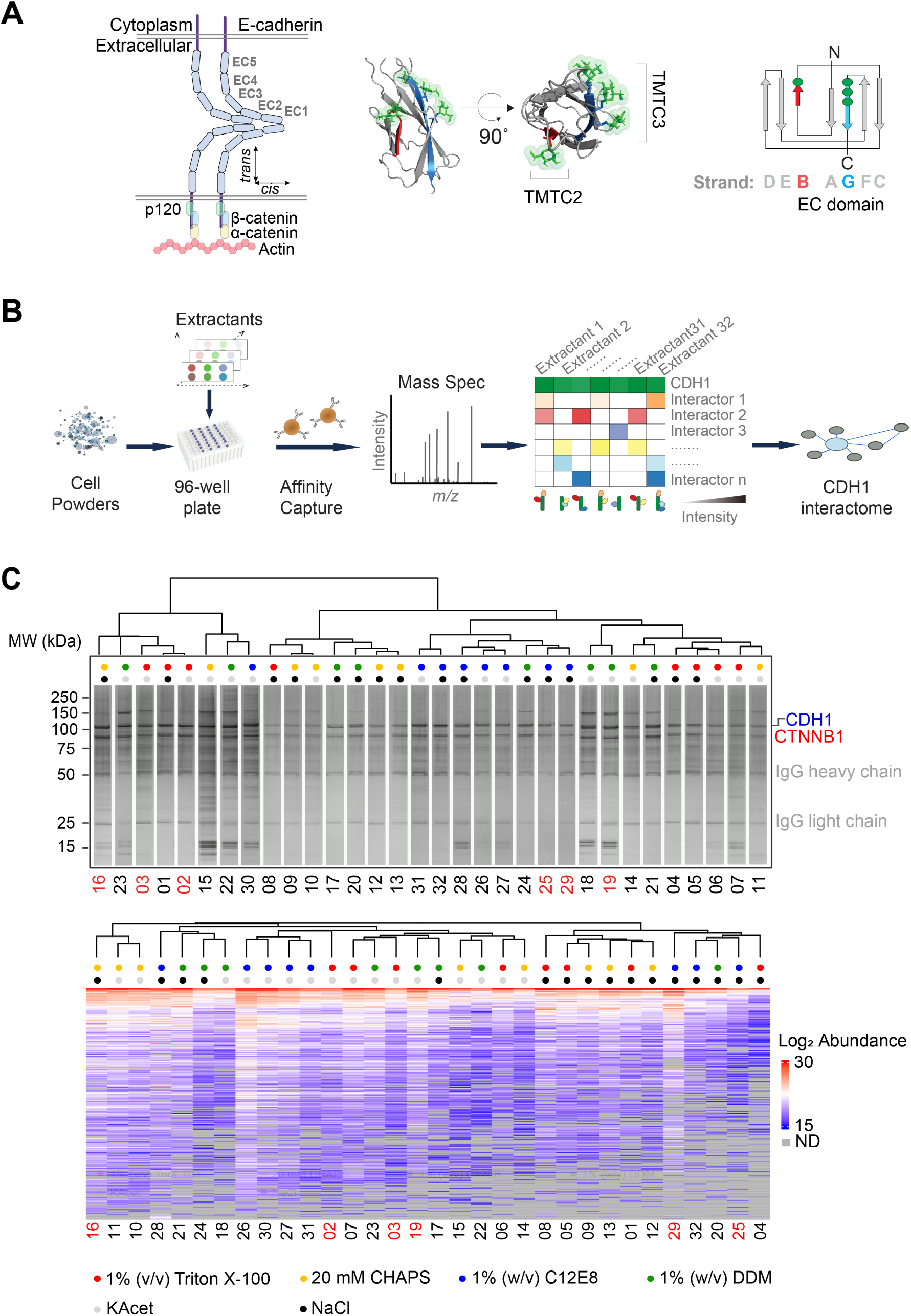
Mapping the *O* -Man dependent CDH1 interactome using IP screening. **(A)** *Schematic diagrams and structural model of CDH1 EC domains*: (left) CDH1 is a transmembrane protein with five EC domains that form cis- and trans-interactions; (middle) TMTC2 mediates *O* -Man on CDH1 EC B-strands, while TMTC3 mediates glycosylations on G-strands (*O* -Man structures were grafted onto an AlphaFold model of EC4 using the GlycoShape tool - mannoses are depicted as green sticks and translucent surfaces on recipient serine and threonine residues (82)); (right) schematic of the β-strand arrangement of an EC domain, highlighting O-Man sites (green dots) on the B-(red) and G-(blue) strands of EC2-4. **(B)** *Schematic diagram of the IP-MS-based interactome screen applied to CDH1* : Cryomilled cells are distributed to a 96-well plate and combined with different extractants; CDH1-associated complexes are affinity enriched from each extract using an antibody coupled magnetic medium and then analyzed by protein MS; the compositions of the enriched macromolecular assemblies will vary according to the stabilizing/destabilizing responses of the protein constituents and a putative interactome is constituted by the combined results. **(C)** *Results of the IP screen using 32 extraction conditions*: (upper) silver-stained SDS-PAGE gel showing CDH1 capture by IP screening; (lower) hierarchical clustering of MS data, with log2-transformed protein abundance values from Proteome Discoverer displayed by color. Grey shading in the heatmap indicates proteins not detected (ND). Six extractants, highlighted in red, were selected for further quantitative analysis. Selected reagents present in <u>extraction solutions are labeled with colored dots.</u>

TMTCs comprise four isoenzymes (TMTC1-4), with different specificities, that catalyze *O* - linked mannose glycosylations (*O* -Man) of cadherins and protocadherins; each TMTC modifies specific Ser/Thr residues, adding single *O* -Man glycans to distinct *β*-strands within particular EC domains of the target proteins (13, 14). Mutations in *TMTC2* and *TMTC4* are associated with nonsyndromic hearing loss (16–20). Mutations in *TMTC3* lead to brain malformations characterized by neuronal migration defects (21, 22), including cobblestone lissencephaly and periventricular nodular heterotopia (PVND). Thus, genetic deficiencies in individual *TMTC* genes underlie distinct neuro-developmental disorders in humans, which suggests that the *O* -Man pattern on EC domains of cadherins and protocadherins serves important roles during organismal development. Yet, the functional roles of these modifications are poorly understood.

Epithelial (E)-cadherin (CDH1) is a widely-studied, classical cadherin that contributes to cell-cell adhesion and, for this, is dependent on five EC domain repeats (EC1-EC5) and a cytoplasmic domain (23). The EC-domains of CDH1 are modified with both *N* - and *O* - glycans, including *O* -Man. With respect to *O* -Man, CDH1 is targeted by TMTC2 (mannose added to EC domain B-strands) and TMTC3 (mannose added to EC domain G-strands) (13, 14, 24); TMTC1 and TMTC4 are predicted to target other cadherin substrates and do not transfer mannose to CDH1 (see *Discussion*). The CDH1 molecules of neighbouring cells engage in homophilic interactions by forming EC1-EC1 dimers, *in trans*. Lateral interactions, involving EC1 and EC2 domains, organize CDH1 into lattices at adherens junctions, *in cis* (25). Furthermore, CDH1 forms functional assemblies with cytosolic *α*-, *β*- and *γ*-catenins shortly after translation, which are necessary for interactions with the actin cytoskeleton in mature adherens junctions (26). These interactions and *O* -Man are depicted in Figure 1A.

Reduced *CDH1* expression is associated with epithelial-to-mesenchymal transition and tumor metastasis (27–29) - contact between fewer CDH1 proteins will reduce cell-cell adhesion potential and promote cancerous cells to escape from the primary tumor site. Previous studies have also suggested that there are alternative or complementary molecular mechanisms by which CDH1 cell adhesion functions may be regulated, independent from altering its absolute expression-level (30); modulating CDH1 PTMs is one such candidate mechanism (31–33). For example, aberrant *N* -glycosylation of CDH1 has been suggested to compromise CDH1 functions (34,35) and more recent evidence indicates that TMTC3 is important for CDH1-mediated cellular adherence (32), suggesting that its loss may negatively impact CDH1 function (i.e., via the loss of EC domain G-strand *O* -Man). Considering the general importance of *N* - and *O* -glycosylations for regulating protein stability, maturation, and functions (36, 37), it stands to reason that *O* -Man modifications on CDH1 EC-domains may impact its functional potential; we therefore sought to interrogate this possibility in the present study.

One way to affect a protein’s functional potential is to alter its ability to participate in physical interaction networks (or ‘interactomes’) (38). Previous studies have investigated the interactome of CDH1 using proximity biotinylation proteomics to identify binding partners of the extracellular or cytoplasmic regions separately (39, 40). However, a comprehensive study of intact CDH1 protein interactions, particularly those that correlate with different *O* -Man states, has not yet been reported. To address this knowledge gap, we conducted comprehensive affinity proteomic analyses of the proteins that co-immunoprecipitate with hemagglutinin (HA)-peptide-tagged, endogenous CDH1 in the presence and absence of *O* -Man glycans. We herein report (i) a CDH1 interactome, obtained from BG1 ovarian cancer cells, and (ii) changes in the protein constituents of CDH1-associated macromolecular assemblies when *O* -Man are altered. Taken together, we survey and dissect macromolecular interactions that may regulate CDH1 functions.

## Methods

### Cell culture

BG1 cells (RRID:CVCL_6570) were cultured adherently in RPMI 1640 medium supplemented with 10% (v/v) FBS. For I-DIRT experiments, cells were cultured using the RPMI 1640 SILAC Protein Quantitation Kit (#A33973, Thermo Scientific). L-proline (200 mg/L; #88211, Thermo Scientific) was added to the medium, which was then filter-sterilized using a 0.22 *µ*m filter (41). Cells were grown in SILAC medium for more than five doublings in a 6-well plate (≳95% incorporation of ^13^C_6_^15^N_4_-Arginine (Arg10) and ^13^C_6_^15^N_2_-Lysine (Lys8) into proteins was confirmed by mass spectrometry) and then expanded to a 500 cm² Culture Dish (#431110, Corning). Cells were harvested and cryomilled as previously described (42).

### Knockout of TMTCs in BG1 Cells

BG1 cells expressing C-terminally HA-tagged CDH1 (BG1*^CDH1::HA^*) were generously provided by Dr. Francis Jacob’s group (University Basel) (43). The following procedure was used to knockout (KO) *TMTCs* from BG1*^CDH1::HA^*. For transfection, BG1*^CDH1::HA^* cells were seeded in a single well of a 6-well plate one day prior to reaching approximately 50% confluence. A DNA mixture (1.0 *µ*g gRNA(s) plasmid + 1.0 *µ*g pBKS plasmid) was combined with 200 *µ*L of OptiPRO SFM Medium (#12309050, Thermo Fisher Scientific) by gentle pipetting (Supp. Table 1). Subsequently, 4 *µ*L of FectoPRO reagent was added to the diluted DNA and mixed thoroughly by gentle pipetting. The cell culture medium was aspirated, and cells were washed once with warm PBS before adding 500 *µ*L of OptiPRO™ SFM Medium to each well. The transfection mixture was incubated at room temperature for 15 minutes before being added to the cells. After two hours, 2 *µ*L of FectoPRO Booster reagent was added to the cells along with 3 mL of RPMI 1640 medium. FACS enrichment of GFP-positive cells was performed 2 days after transfection. The enriched cells were cultured under normal conditions for an additional week before being single-cell sorted into a 96-well plate (44, 45). KO clones were screened using Indel Detection by Amplicon Analysis (IDAA) and further confirmed by Sanger sequencing, as previously described (Supp. Table 2) (44).

### IP pre-screens

The experimental schema is depicted in Figure 1B (as previously described (46–48). Extractants containing different detergents were initially pre-screened (pre-screen #1) for their effects on CDH1 yield by IP (Supp. Table 3); this was followed by another prescreen (pre-screen #2), inspired by a fractional factorial design (49), of 32 unique extractant formulations (listed in Supp. Table 4) to discover suitable I-DIRT screening conditions (described below) from BG1*^CDH1::HA^* cells. Briefly, ∼37.5 mg of cell powder was dispensed into wells of a 96-well plate using spacer (#187) and extracted in 500 *µ*L of extraction solution with 1 protease inhibitor cocktail (#11836170001, Roche). Samples were sonicated using a QSonica Q700 with an 8-tip microprobe (#4602, QSonica) at an amplitude of 1 for 30–40 s continuously at 4*^◦^*C. Extracts were transferred to 1.5 mL microtubes using an 8-channel pipette and centrifuged at 20,000 g for 10 min at 4*^◦^*C. Clarified supernatants were transferred to a new 96-well plate containing 5 *µ*L of epoxy beads (#14301, Thermo Scientific) conjugated with the *α*-HA antibody (#26183, Thermo Scientific). Samples were incubated for 30 min at 4*^◦^*C with gentle mixing to allow for affinity capture. Beads were washed with a detergent-free solution that was otherwise identical in composition to the initial extractant. Captured proteins were eluted with 50 *µ*L of 2% (w/v) SDS in 40 mM Tris-HCl for 5 min at 70*^◦^*C with vigorous shaking. 10 *µ*L of each eluate (20% of the total volume) were used for SDS-PAGE with silver staining kit (#10543053, Thermo Scientific; Fig. 1C). Stained gel images are also available in an interactive, searchable format at http://www.copurification.org (navigate to “search public gels,” select species: human, tagged protein: CDH1, then click “next” and follow prompts). The remaining 80% of each eluate was processed using S-Trap™ columns for MS; raw data were analyzed using Proteome Discoverer (v2.4). To compare the IP results from general protein staining with those from LC-MS/MS, we performed hierarchical clustering on the quantified protein abundances. 6 of the 32 extractants were advanced to the next round, for I-DIRT-based quantitative interaction screening, in quadruplicates (see below).

### I-DIRT IP screen

Selected 6 of the 32 extractants from IP screening were moved forward into I-DIRT to assess the specificity of observed interactions: quadruplicate IP replicates by I-DIRT (50–52), using 24 wells of a 96-well plate for forward (heavy-labeled BG1*^CDH1::HA^* mixed with light-labeled BG1^WT^) and reversed (light-labeled BG1*^CDH1::HA^* mixed with heavy-labeled BG1^WT^) screening, respectively (see also: *Cell culture*, above). In I-DIRT screening, ∼37.5 mg of 1:1 (w:w) mixed heavy- and light-labeled cell powders were used per well (Supp. Fig. 1). See also: *MS data processing for I-DIRT*, below.

### Label-free IP screen (with TMTC KOs)

Selected 6 of the 32 extractants from IP screening were applied to IP for wild and TMTCs deficient BG1*^CDH1::HA^* cell lines respectively, including BG1*^CDH1::HA^*, BG1*^CDH1::HA/^* ^KO*:TMTC2*^, BG1*^CDH1::HA/^* ^KO*:TMTC3*^, BG1*^CDH1::HA/KO:TMTC2/3^*, and BG1*^CDH1::HA/KO:TMTC1-4^*. For each cell line, 24 IPs were performed in a 96-well plate (quadruplicate IPs for 6 extractants) using approximately 37.5 mg of cell powder per well. See also: *IP pre-screens,* above, and *MS data processing for label-free IP experiments*, below.

### Bulk protein extraction from cells

Approximately 20 mg of cryo-milled cell powders (or mixed powder, as described above) was extracted in 200 *µ*L of 2% (w/v) SDS, 40 mM Tris-HCl containing 1× protease inhibitor cocktail (#11836170001, Roche), followed by 1 min of sonication (3 s on and 3 s off, 35 Amp) at 4*^◦^*C. The extract was centrifuged at 20,000 g for 10 min, and the supernatant was collected. The protein concentration of the clarified extract was measured using BCA assay (#23227, Pierce).

### Protein digestion and peptide sample preparation for LC-MS/MS analysis

Samples were processed using S-Trap™ micro spin columns (#C02-micro-80, ProtiFi) following a modified version of the manufacturer’s micro S-Trap protocol (https://protifi.com/pages/protocols). The normal procedure was used for clarified cell extracts and the ‘high recovery’ procedure was used for IP samples. IP elutions and clarified cell extracts were dried (speed-vac): clarified extracts were redissolved in 23 *µ*L of 5% (w/v) SDS, 50 mM TEAB (pH 8.5); IP eluates were redissolved 23 *µ*L of 5% (w/v) SDS, 8 M urea, 100 mM glycine (pH 7.55); after which the manufacturer’s instructions were followed.

### LC-MS/MS data collection from peptide mixtures

The liquid chromatography tandem mass spectrometry consisted of UltiMate 3000 RSLCnano UHPLC system interfaced with an Orbitrap Exploris 480 mass spectrometer (Thermo Fisher Scientific) with a nanoelectrospray ion source (Thermo Fisher). Briefly, for label-free and I-DIRT IP samples, the peptides eluted from S-traps were dried and resuspended in 25 *µ*L of a water : methanol : formic acid solution (94.9 : 5.0 : 0.1, parts by volume). Of this, 5 *µ*L was loaded onto a 75 *µ*m × 50 cm Acclaim PepMap RSLC nano Viper column filled with 2 *µ*m C18 particles (Thermo Fisher Scientific) over a 60 min elution gradient (Solvent A is 0.1% (v/v) formic acid in water and Solvent B is 0.1% (v/v) formic acid in acetonitrile) with flow at 300 nL/min. The column temperature was set at 40*^◦^*C. The gradient was set as: 3% B over 3 min; 3 to 50% B over 45 min; 2 min to 80% B; then wash at 80% B over 5 min, 80 to 3% B over 2 min and then the column was equilibrated with 3% B for 3 minutes. For a full mass scan, a profile spectrum was acquired m/z 375 to 1,500 at a 120,000 resolution. For the MS2 scan, the top 25 most intense ions with charge +2-6 were selected for fragmentation by HCD with an isolation window of 1.4 m/z units. The normalized collision energy was set to 30%. The exclusion time for previously selected precursors was set as 20 s. The fragment spectra were acquired in centroid mode with a 300% normalized AGC and 50 ms maximum injection time.

For bulk proteins extracted from I-DIRT cell powder mixtures, peptides were dissolved at 500 ng/*µ*L with a water : methanol : formic acid solution (94.9 : 5.0 : 0.1 by volume) and 1 *µ*g of samples were loaded to column over a 150 min elution time. The gradient was set as: 3% B over 3 minutes; 3 to 50% B over 120 minutes; 2 minutes to 80% B; then wash at 80% B over 15 minutes, 80 to 3% B over 2 minutes and then the column was equilibrated with 3% B for 3 minutes. The other settings were the same as for IP samples.

For bulk proteins extracted during label-free analysis of *TMTCs* KOs (including BG1*^WT^*, BG1*^CDH1::HA^*, BG1*^CDH1::HA/^* ^KO*:TMTC2*^, BG1*^CDH1::HA/^* ^KO*:TMTC3*^, BG1*^CDH1::HA/KO:TMTC2/3^*, and BG1*^CDH1::HA/KO:TMTC1-4^*), experiments were performed on Thermo Scientific Orbitrap Exploris 480 with data-independent acquisition (DIA) mode (see also: *MS data processing for Data Independent Acquisition*, below). 1 *µ*g of peptides were loaded to column over a 120 min elution time. The gradient was set as: 2% B over 3 minutes; 3 to 45% B over 87 minutes; 5 minutes to 80% B; then wash at 80% B over 8 minutes, 80 to 2% B over 7 minutes and then the column was equilibrated with 2% B for 10 minutes. MS spectra were acquired in the m/z 400-1200 range at a resolution of 120,000. DIA windows were set to 92 scan events with isolation windows of 8.6 m/z. Fragmentation was performed using 32% normalized collision energy, and MS/MS spectra were acquired at a resolution of 30,000. The normalized AGC target was set to 1,000% with an auto maximum injection.

### MS data processing for I-DIRT

The I-DIRT data was processed using Proteome Discoverer 2.4 (Thermo Fisher Scientific) and searched against the human UniProt database (proteome ID: UP000005640, reviewed; downloaded in August 2020). The quantification method was selected as SILAC (Arg10 and Lys8). All other parameters were set the same as for the label-free data (see also: *MS data processing for label-free IP experiments*, below). Postprocessing was performed with in-house R scripts. Peptides identified in fewer than two replicates of the BG1*^CDH1::HA^* sample were not considered for further analysis. Imputation was applied to the remaining data to compensate for partial missing values, modified from previously described methods (47), as follows. For peptides with missing values in more than two replicates in BG1^WT^, method 3.1 was used: briefly, missing peptide values were imputed by random sampling from a uniform distribution between *µ* - 3*σ* and *µ* - 2*σ*, where *µ* and *σ* represent the mean and standard deviation, respectively, of all protein intensities within the replicate.

For the remaining partial missing peptide values in BG1*^CDH1::HA^* or BG1*^WT^* samples, imputation was performed using method 4.1. This method is based on the distribution of relative differences between observed replicate measurements. For each peptide, a relative difference (Δ) was calculated between observed replicate intensities:

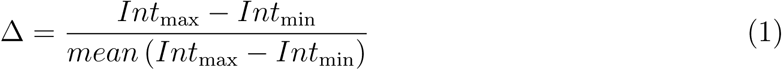

where Int_max_ and Int_min_ are the highest and lowest peptide intensities across replicates. Then new standard deviation of this distribution (Δ*_new_*) was scaled by the average correlation mean(corr) between observed replicate intensities and the intensities of replicates with non-zero values:

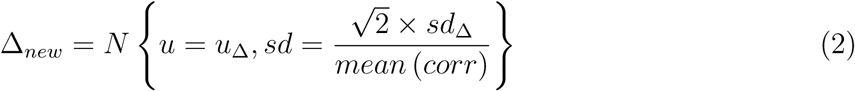

here *N* {µ*, σ*} represents a normal distribution. Finally, missing intensities were imputed as the mean of the observed non-zero replicate intensities, scaled by the absolute value of Δ*_new_*:

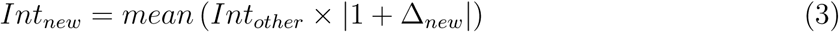

where Int_other_ represents the intensities of the protein in replicates with non-zero values. Then the ratio for I-DIRT was calculated as using the peptide abundance:

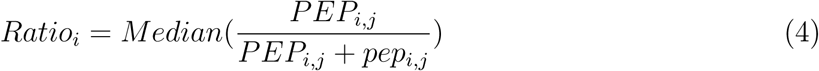

where Ratio_i_ represents the I-DIRT ratio of protein i, PEP_i,j_ represents the abundance of unique peptide j of protein i in BG1*^CDH1::HA^*, pep_i,j_ represents the abundance of unique peptide *j* of protein *i* in BG1^WT^.

Two criteria were implemented to filter for statistically significant specific interactors from I-DIRT experiments: (i) a Bayesian factor greater than 3, and (ii) location within a validation ellipse in the I-DIRT versus swap ratio plot. For the Bayesian test, the ratios for each protein in I-DIRT/swap replicates (the IPs) were compared against the general median ratios of all proteins in the clarified cell extract used for the IPs (starting material obtained from mixed heavy- and light-labeled cell powders). The ellipse was centered at (1,1), with its semi-major and semi-minor axes determined by the median and standard deviation (SD) of bulk protein ratios from the original and swapped cell powder mixes. Only interactor candidates that passed the Bayesian factor analysis within the ellipse were considered as specific CDH1 interactors.

### MS data processing for label-free IP experiments

LC-MS/MS raw data from IP experiments, acquired using Data Dependent Acquisition (DDA), were processed using Proteome Discoverer 2.4 (Thermo Fisher Scientific) and searched against the human UniProt database (proteome ID: UP000005640, downloaded in August 2020). Full trypsin specificity was set with up to two missed cleavage sites allowed. Fixed modification of MMTS on cysteine residues (+45.988 Da) was specified, along with variable modifications including methionine oxidation (+15.995 Da), protein N-terminal acetylation (+42.011 Da), Met-loss (−131.04 Da), and Met-loss+Acetyl (−89.030 Da). Precursor and fragment tolerances were set to 10 ppm and 0.02 Da, respectively. Identification results were filtered with 1% FDR. Data was post-processed using in-house R scripts. Only proteins identified with at least two unique peptides were considered. Protein abundance values (normalize intensities) were log_2_-transformed. Missing values were imputed using the same approach as for I-DIRT data (above), but this was done at the protein abundance level. All protein abundances were normalized to CDH1 to enable direct comparisons. Unpaired, two-sample t-tests were performed to compare protein abundances between *TMTCs* deficient BG1*^CDH1::HA^* and BG1*^CDH1::HA^* in IP experiments. The significantly perturbed interactors were filtered by log_2_ fold change ≥ 1 or ≤ −1 with p-value ≤ 0.05.

### MS data processing for Data Independent Acquisition

DIA HRMS1 data were processed using DIA-NN (v 1.8.1) in library-free mode (53, 54). The following modifications were applied: fixed modification of MMTS on cysteine residues (+45.988 Da), and variable modifications including methionine oxidation (+15.995 Da) and protein N-terminal acetylation (+42.011 Da). The precursor FDR was set to 1%. All other parameters were left at their default. Post-processing was performed with FragPipe-Analyst (http://fragpipe-analyst.nesvilab.org/).

### Differential O-Man glycoproteomics

Sample preparation for differential glycoproteomics was as previously described with minor modifications (55). Briefly, BG1^WT^ and BG1*^CD1::HA/KO: TMTC1-4^* cell pellets were extracted with 0.5% (w/v) SDS, 40 mM Tris, pH 8.0 on ice followed by reduction (10 mM DTT, 72*^◦^*C, 10 min) and alkylation (25 mM IAA, RT, 20 min). Triton X-100 was added to 1% (v/v) before PNGase F digestion (8U, 37*^◦^*C, 16h) and trypsin digestion (25 *µ*g, 37*^◦^*C, 16h). Tryptic digests were desalted by Sep-Pak C18 (100 mg) cartridges and labeled by diethyl stable isotopes. Glycopeptides were enriched by BC2L-A lectin chromatography, analyzed by Orbitrap Fusion mass spectrometry (2h gradient, HCD/ETciD fragmentation) and identified/quantified with Proteome Discoverer 1.4.

### Cell adhesion assay

Cell adhesion was measured using the Vybrant™ Cell Adhesion Assay Kit (V13181, Thermo Fisher Scientific). Briefly, different cell lines were dissociated with EDTA and labeled with calcein AM. 2 × 10^5^ cells were then seeded onto a monolayer of BG1^WT^ cells in a 96-well plate. The assay was performed according to the manufacturer’s instructions.

### Flow cytometry

Cells were grown in 6 wells and at 80% confluency they were washed with PBS and detached from cell culture dishes using EDTA based reagent CellStripper for 15 min. Detached cells were washed with PBS once and 50 000 cells were distributed in round bottom 96-well plates and kept on ice throughout the experiment. Anti-CDH3 (MAB861, R&D systems 1:500 in 10% (v/v) FBS in PBS) and Anti-CDH1 primary antibodies (AF648 R&D systems 1:200 in 10% (v/v) FBS in PBS) were added to the cells and incubated for 1 h. The cells were washed 3 times with 100 ul of 10% (v/v) FBS in PBS and incubated with secondary antibody conjugated to fluorophore (Goat anti-Mouse IgG (H+L) Cross-Adsorbed Secondary Antibody, Alexa Fluor™ 488 or Donkey anti-Goat IgG (H+L) Cross-Adsorbed Secondary Antibody, Alexa Fluor™ 488 at 1ug/ml in 10% (v/v) FBS in PBS). The analysis was performed using the Beckman Coulter Life Sciences CytoFLEX flow cytometer.

### Immunofluorescence

Cells were seeded in 24 well plates onto poly-L-lysine precoated coverslips. Two days later the cells were washed with PBS and fixed in 4% PFA for 10 min at RT. Fixed cells were washed 3 times in PBS and stored at 4*^◦^*C until staining. Before staining, cells were permeabilized with 0.3% Triton X-100 in PBS for 10 min and rinsed 3× 10 min in PBS containing 0.3% (w/v) glycine. Blocking of nonspecific binding was achieved by incubating the cells in 5% (w/v) BSA in PBS, 0.3% (w/v) glycine. Cells were incubated in anti-CDH3 primary antibodies (C13F9, Cell Signalling, 1:50 and MAB861, R&D systems 1:100 in 5% [w/v] BSA in PBS, 0.3% [w/v] glycine) and anti-HA high affinity primary antibody (3F10, Roche 1:200 in 5% [w/v] BSA in PBS, 0.3% [w/v] glycine) rinsed 3x in PBS for 10 min each and incubated in diluted conjugated secondary antibodies (goat anti-rabbit IgG, goat anti-mouse IgG Alexa Fluor™ 594 as well as Donkey anti-Rat IgG (H+L) Cross-Adsorbed Secondary Antibody, Alexa Fluor™ 488 1:500 in 5% [w/v] BSA in PBS, 0.3% [w/v] glycine) and DAPI [1:1000]). After washing in PBS, the coverslips were mounted to glass slides using Prolong Gold mounting media. Zeiss LSM900 high-resolution confocal microscope was used for imaging.

### Western blotting

Briefly, cells were grown for 2 days until 80% confluency, washed in PBS and then lysed with 50 mM Hepes pH 7.4, 150 mM NaCl, 1% (v/v) Triton-X100 and Roche Complete protease inhibitor mix. After 30 minutes on ice, the lysate was cleared by centrifugation (10,000 g, 10 min), normalized using BCA assay (Thermo), diluted to 1x LDS (Invitrogen), reduced with dithiothreitol (15 min, 60*^◦^*C) and run on Bis-Tris 4-12% 10-well precast gels (Invitrogen) in 1x MES buffer (Invitrogen). Proteins were transferred (1h, 100V) to ethanol-activated PVDF membrane using Tris-Glycine transfer buffer (25 mM Tris, 192 mM glycine, 10% EtOH). Membrane was incubated for 1h at RT in agitation with primary monoclonal antibodies Anti CDH3 (MAB861, R&D systems 1:500 in blocking milk). Secondary anti-mouse HRP-conjugated secondary antibody (Dako, 1:4000 in blocking milk) was used for the CDH3 detection, after washing in TBST. For *β*-actin staining, a parallel blotting was performed using *β*-actin conjugated primary antibody (Abcam ab49900; 1:25000 in blocking milk). HRP signal was obtained using SuperSignal™ West Pico PLUS Chemiluminescent Substrate (Thermo Scientific) and acquired using GE Healthcare ImageQuant™ LAS 4000 imager.

## Results

### Construction of cell models for capturing CDH1 interactomes +/− *O* -Man

To ensure our studies could be conducted effectively using a well-validated, commercially available antibody to target our protein of interest, we elected to work with epitope-tagged CDH1 protein. *CDH1* is expressed in the near-diploid human ovarian adenocarcinoma cell line, BG1 (56); and conveniently, a BG1-derived cell line expressing C-terminally HA-tagged CDH1 from the native loci (CRISPR/Cas9-based knock-in, BG1*^CDH1::HA^*) was available (43, 45). We therefore elected to work with these cells and further engineered them using a CRISPR/Cas9-based knock-out (KO) strategy, to establish individual and combinatorial genetic ablation of *TMTCs* including single KO of *TMTC2* (BG1*^CDH1::HA/KO: TMTC2^*) and *TMTC3* (BG1*^CDH1::HA/KO: TMTC3^*), double KOs of *TMTC2/3* (BG1*^CDH1::HA/KO: TMTC2/3^*), and quadruple KO of *TMTC1-4* (BG1*^CDH1::HA/KO: TMTC1-4^*); see Supplemental Tables 1 & 2. This collection of cell lines allowed us to interrogate CDH1 interactomes in different *O* -Man states using an immunoprecipitation and mass spectrometry-based (IP-MS) interactome screen, as depicted in Figure 1B (46–48).

### Immunoprecipitation optimization for CDH1

In an attempt to comprehensively delineate a CDH1 interactome, we explored a multitude of extractants, intending to reveal ‘concealed assemblies’ that often go missed in unoptimized experimental conditions (46, 48, 57–59). We started by pre-screening reagents, formulating candidate extractants to use for detailed IP-MS analyses of CDH1-associated macromolecular assemblies, present within BG1*^CDH1::HA^* cells. Pre-screening is a quality assurance practice, ultimately allowing the judicious selection of extractants that exhibit features conducive to bona fide interaction discovery. As a membrane protein, we reasoned that CDH1 will require special attention to the selection of the detergent components of the extractants used during co-IP; membrane proteins, like CDH1, which may rely upon the presence of residual lipid membrane fragments and/or detergent surrogates, originating from the extractant, to remain soluble and/or maintain authentic associations (46, 59–63). Multiple detergents were evaluated and the results demonstrated a significant influence on CDH1 extraction efficiency; for example, extractants containing Tween 20 or Brij-58 exhibited reduced CDH1 yields (Supp. Table 3).

Taken together, these results informed our creation of an extractant pre-screen consisting of thirty-two unique formulations (Fig. 1C & Supp. Table 4), comprising five distinct factors, each with variations: (i) pH buffer type (HEPES-Na or Tris-HCl, each at a set concentration and pH), (ii) detergent type (Triton X-100, CHAPS, DDM, or C12E8, each at a set concentration), (iii) salt type (NaCl or CH_3_COOK) and (iv) its concentration (250 or 500 mM), and (v) calcium (added at 1 mM, or not added). Protein profiles from the IP pre-screen were initially visualized using SDS-PAGE with silver staining. The lanes displayed a few consistently dominant protein bands (Fig. 1C): CDH1 was captured with some variations in enrichment efficiency displayed across extractants; a band at ∼90 kDa (CTNNB1 is a known CDH1 interactor whose yield is closely correlated with the CDH1 yield; Supp. Fig. 2A); bands observed at ∼50 and ∼25 kDa (attributable to IgG chains that are leached from the affinity medium upon elution in a hot SDS-containing solution (2% [w/v] at 70*^◦^*C)); and we noticed that IPs from potassium acetate-containing extractants tended towards higher proteinaceous complexity and the accumulation of a trio of bands at ∼15 kDa (see extractants 15, 22, 30, 18, 19, and 07). Condition-specific variations in the apparent recovery of many low intensity bands was also apparent.

While the gel banding pattern gives us a sense of how the protein co-IP landscape is influenced by the extractant, it provides no explicit insight into the underlying variety of protein molecular identities and functions at play. Hence, the same fractions were analyzed by LC-MS/MS and the results were visualized as a hierarchical protein abundance heatmap (Fig. 1C). This readout provides us with an estimation of the ‘interactomic space’ explored by the pre-screen, *i.e.*, how similar or different the results produced by each extractant are at the protein-ID level. Weighing our visual inspection of the stained gel with the semi-quantitative MS analysis, we selected six conditions (Fig. 1C) to advance for screening by quantitative MS analysis (below). These extractants encompassed over 95% of proteins detected across all thirty-two initial extractants (Supp. Fig. 2B; and surpassed the coverage of 85% of one thousand randomly selected six-extractant combinations, Supp. Fig. 2C). Correlation analysis revealed high intra-group similarity (e.g., extractants 3 versus 16/25; 19 versus 29) contrasted with significant inter-group heterogeneity (e.g., 19/29 versus 2/3/16/25; Supp. Fig. 2D & E).

### CDH1 interactome

To accurately discern which candidate CDH1 interactors were formed within the BG1 cells, we employed the isotopic <u>d</u>ifferentiation of interactions as <u>r</u>andom or <u>t</u>argeted (I-DIRT) technique (50–52), with four replicates of each selected condition. The label-swap I-DIRT design is depicted in Fig. 2A. BG1*^CDH1::HA^* and BG1^WT^ cells were cultured in the presence of media containing heavy (Lys8; Arg10) or naturally occurring carbon and nitrogen isotopes within lysine and arginine (*i.e.*, SILAC; see *Methods*). Isotope-labeled BG1*^CDH1::HA^* and BG1^WT^cell powders were mixed in equal amounts as the starting materials for I-DIRT/swap experiments (Supp. Fig. 1). This approach enables the classification of protein interactions by IP-MS according to the following categories: (i) constituents formed in physical assemblies along with CDH1 (the IP target), *in vivo* (in the cell), that also remain stably associated with CDH1, *in vitro* (in the test tube, post-extraction), appear in IP-MS data predominantly as heavy-isotope labeled peptides (e.g., true positives are typically ∼80-100% heavy); (ii) non-specific, post-extraction artifacts (noise) and bona fide *in vivo*interactors that are unstable or dynamic *in vitro* (false negatives), equilibrate on the time-scale of the IP, and will converge on equal heavy- and light-isotope labeling (distributed around ∼50% +/− ∼10% heavy); and (iii) real *in vivo* interactors that exhibit intermediate stability *in vitro* will not fully equilibrate on the time scale of the IP and will converge on intermediate heavy isotope labeling (e.g., between ∼60-80% heavy (50–52). The above-described *in vitro* behaviors will be influenced by the extractant used (see *Discussion*).

**Figure 2.**
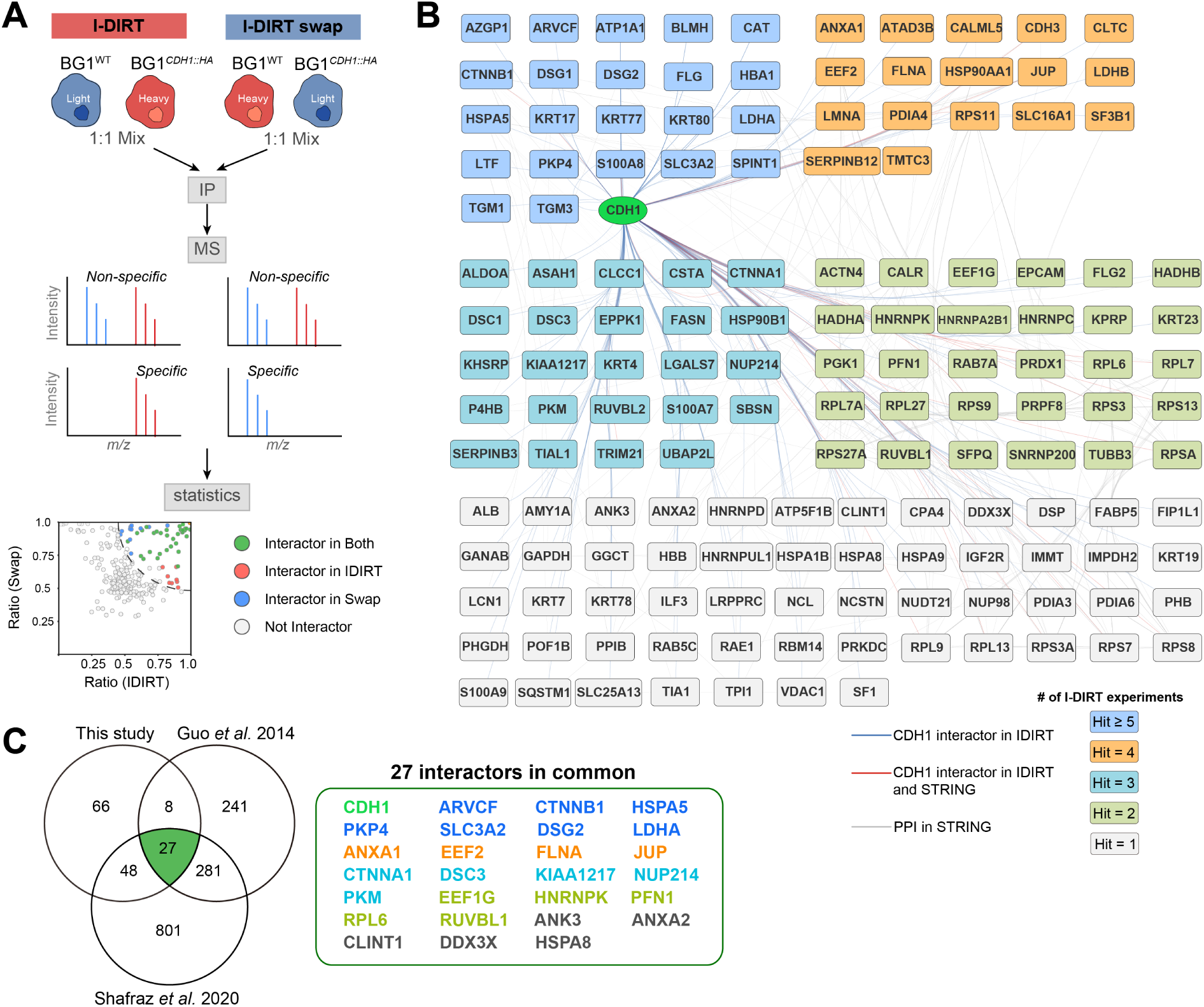
I-DIRT screen of CDH1. **(A)** *Depiction of the approach*: BG1^WT^ cells and BG1*^CDH1::HA^* cells were cultured in both light and heavy isotope-labeling media for label-swapped I-DIRT experiments. The resulting cell powders were combined in 1:1 (w:w) ratios for IP-MS analyses. Specific CDH1 interactors are enriched in one isotope-labeled channel in MS, while non-specific interactors are quantified comparably in both heavy and light channels. BG1*^CDH1::HA^* cells cultured in heavy-isotope media were designated ‘I-DIRT,’ while BG1*^CDH1::HA^* cells cultured in light-isotope media were designated ‘I-DIRT swap.’ **(B)** *Specific interactors identified across six I-DIRT experimental conditions*: the interactors are grouped based on how many times they were identified as specific interactors. Blue lines connect CDH1 interactions identified in this study, orange lines connect CDH1 interactions also annotated in the STRING database (83), while gray lines connect other protein interactions annotated in the STRING database that are supported by these data. **(C)** *Overlap of the I-DIRT interactor list with two previously published datasets* (39, 40): the panel on the right lists the 27 common interactors, colored by their identification frequency, as in (B).

While traditional proteomics approaches typically rely on protein-level quantification, we observed that in these I-DIRT experiments, protein quantitative measurements were sometimes biased by aberrant quantitation of either heavy- or light-labeled peptides. For example, CDH1 and its established interactor CTNNB1 exhibited more consistent I-DIRT ratios using peptide-level rather than protein-level calculations (Supp. Fig. 3). The importance of peptide-level analysis was further exemplified by the case of DSG1, where protein-level quantification yielded discordant ratios between label-swapped I-DIRT experiments, while peptide-level analysis maintained consistency across experimental conditions (Supp. Fig. 4). Detailed examination of DSG1 peptides revealed that a single outlier peptide exhibited 1000-fold higher abundance than others and significantly distorted the protein-level quantitation (Supp. Fig. 5). These observations suggested that peptide-based quantitation is more robust and we consequently used median peptide ratios for our calculations (see *Methods*).

Specific interactors identified in I-DIRT experiments for each extractant were pooled to define the CDH1 interactome, consisting of 148 hits (Fig. 2B & Supp. Fig. 6; interactive plots are available at https://ngrc-glycan.shinyapps.io/CDH1_interactome/). Fourteen proteins passed our filters (see *Methods*) for specific CDH1 interactors under all six IP conditions (including ARVCF, AZGP1, CAT, CTNNB1, DSG1, HBA1, HSPA5, KRT77, LTF, PKP4, S100A8, SLC3A2, SPINT1, TGM3), while others, such as PDIA3, and PDIA6 were identified only under certain extractants (extractant 29 and extractant 2, respectively). The distribution of specific interactors identified in each extractant is presented in Supplemental Figure 6. We compared the CDH1 interactome identified in this study with two previously published datasets that used proximity labeling methods; a total of 27 common interactors were found across all three datasets (Fig. 2C). Gene Ontology and Reactome pathway analyses (Fig. 3A-D) revealed significant enrichment in cell adhesion-related categories (cell-substrate junctions, focal adhesions) and epidermal development pathways (skin development, epidermal differentiation), as well as dynamic processes (cell-cell communication, neutrophil degradation). These findings suggest the co-enriched proteins cooperate with CDH1 to mediate both structural and regulatory functions in epithelial biology. Manual curation using UniProt GO annotations identified 44 junction, plasma membrane, or secreted proteins among the E-cadherin interactors, such as CDH3, PDIA6, ANXA1, and ANXA2 (Fig. 3E; (64)).

**Figure 3.**
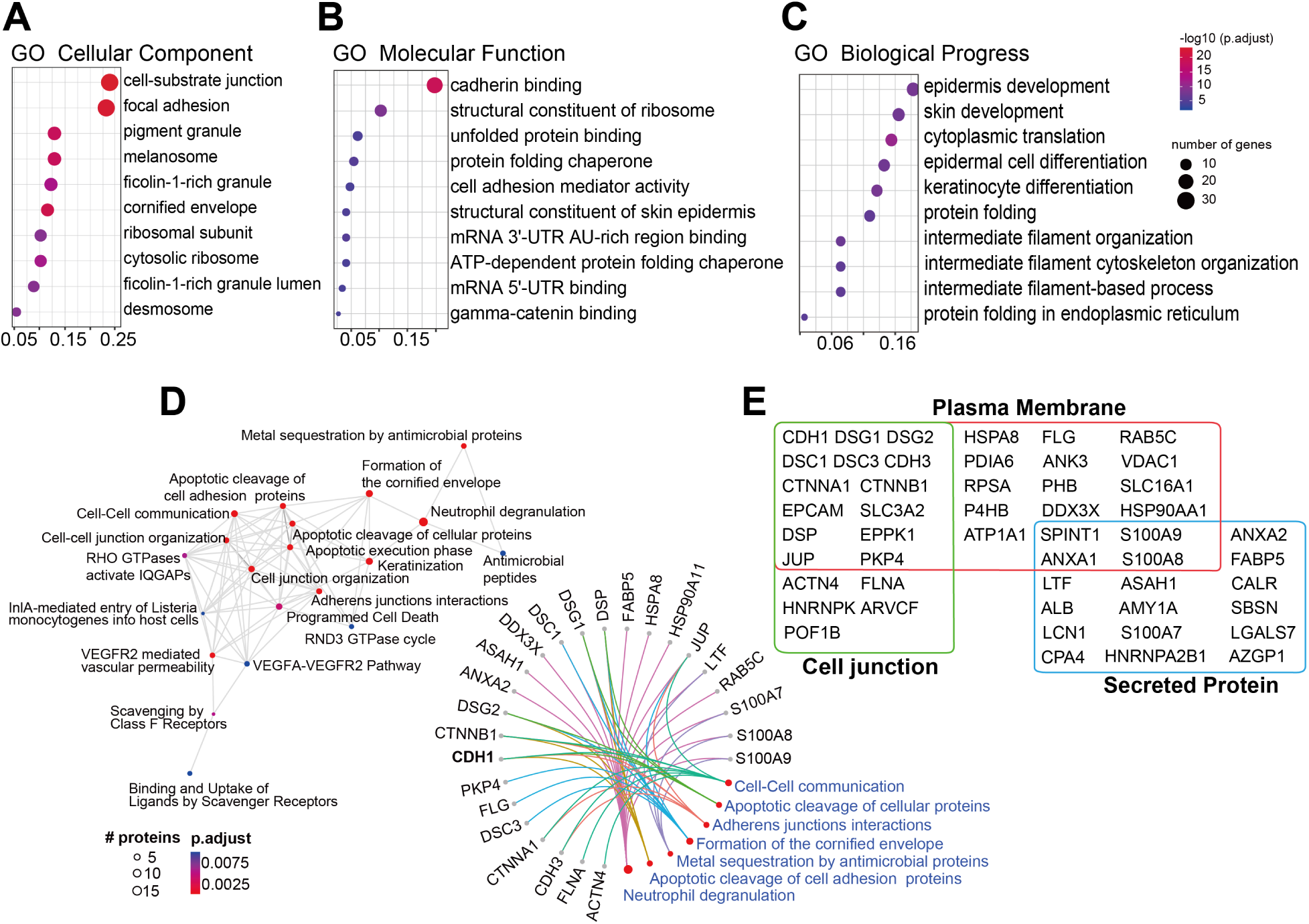
Bioinformatic analyses of CDH1 interactors. *Gene Ontologies* (GO) **(A–C)** and *Reactome pathways* **(D)** enriched among the specific CDH1 interactors. **(A)**Enrichment of GO cellular components (CC) associated with CDH1 interactors. **(B)** Enrichment of GO Biological Processes (BP) associated with CDH1 interactors. **(C)** Enrichment of GO Molecular Functions (MF) associated with CDH1 interactors. **(D)** The enriched Reactome pathways are shown on the left. The specific CDH1 interactors involved in each pathway are detailed on the right. **(E)** *Localizations of select CDH1 interactors* : proteins with annotated localizations at the cell surface or extracellular matrix are listed.

### The *O* -Man dependent CDH1 interactome

The recently discovered TMTC-driven *O* -Man pathway plays a crucial but poorly understood role for cellular functions. Under the assumption that *O* -Man on different structural components (i.e., G-strand versus B-strand) of EC domains could differentially impact protein interactions formed with CDH1, we systematically investigated the *O* -Man-dependent CDH1 interactome. For this, we employed multiple *TMTC* KO cell lines, constructed in the BG1*^CDH1::HA^* genetic background, including BG1*^CDH1::HA/KO: TMTC2^* (*O* -Man depletion on EC B-strands), BG1*^CDH1::HA/KO: TMTC3^* (*O* -Man depletion on EC G-strands), as well as BG1*^CDH1::HA/KO: TMTC2/3^* (encompasses complete CDH1 *O* -Man depletion), and BG1*^CDH1::HA/KO: TMTC1-4^* (encompasses complete cadherin / protocadherin *O* -Man depletion). We first verified that *O* -Man is depleted in our BG1 *TMTC1-4* KO (BG1*^CDH1::HA/KO:TMTC1-4^*) cell line through diethyl stable isotope (DEL) analysis, as previously described (Supp. Fig. 7; (55)). The analysis revealed selective loss of *O* -Man glycans on CDH1 and CDH3, while *O* -Man remained unchanged on POMT1/2 substrates (*α*-dystroglycan and SUCO) and on TMEM260 substrates (c-MET and RON) in *TMTC1-4* KO cells, as previously described in HEK293 cell models (55). We then performed differential interaction profiling by (label free) IP-MS, statistically comparing proteins co-isolated with HA-tagged CDH1 in IPs from *TMTC* KO cells to those from BG1*^CDH1::HA^* cells. The profiling was done using the same six IP conditions as in I-DIRT, each in quadruplicates. Only proteins that were likely to have formed interactions with CDH1 in cells, as assessed by I-DIRT, were considered for further analysis. The results are summarized in Figure 4; a comprehensive overview is provided in Supplemental Figure 8 and can also be explored using our interactive web resource: https://ngrc-glycan.shinyapps.io/CDH1_interactome/. To visualize the *O* -Man-mediated behavioral similarities exhibited by CDH1 interactors, we performed hierarchical clustering of the average abundance ratios of the interactors (from six IP conditions) in the four *TMTC* -deficient cell lines, relative to those in BG1*^CDH1::HA^*. Figure 4A summarizes the behaviors of different CDH1 interactors upon *TMTC* KO: five main clusters display the direction of interaction changes with CDH1 (increased or decreased co-enrichment) for each I-DIRT specific protein. Group 1 includes interactors whose co-enrichment with CDH1 were consistently decreased, using any of the *TMTC* KO cell lines, in at least one IP condition (the median of log_2_ fold change values for all Group 1 interactors in *CDH1::HA* IPs from *TMTC2*, *TMTC3*, *TMTC2/3*, and *TMTC1-4* KO cells, compared to *TMTC* WT, were −3.4, −3.5, −3.7, −2.6, respectively). Group 2 includes interactors whose co-enrichment with CDH1 was more mildly decreased by *TMTC* KOs, with diverse behaviors observed under conditions of *O* -Man glycan depletion (for example, DSG2 (Group 2) co-enrichment decreased in fewer tested IP conditions than CAT (Group1), shown in Fig. 4B; the median of log_2_ fold change values for all Group 2 interactors in CDH1::HA IPs from *TMTC2*, *TMTC3*, *TMTC2/3*, and *TMTC1-4* KO cells, compared to *TMTC* WT, were −1.7, −1.8, −2.0, −1.3, respectively). Group 3 contains interactors whose co-enrichment with CDH1 was relatively unaffected by *TMTC* KO status, suggesting their interactions with CDH1 are not physically influenced by *O* -Man. Groups 4 and 5 (orange and red, respectively) represent interactors displaying increased co-enrichment with CDH1 upon *TMTC* KO. The median of log_2_ fold change values for all Group 4 interactors in CDH1::HA IPs from *TMTC2*, *TMTC3*, *TMTC2/3*, and *TMTC1-4* KO cells, compared to *TMTC* WT, were −0.2, 0.2, −0,1, 1.0, respectively; a mild co-enrichment largely dominated by effects observed in *TMTC1/4* KO. The median of log_2_ fold change values for all Group 5 interactors in four *TMTCs* KO cell lines were 1.4, 1.7, 1.5, 2.4, respectively.

**Figure 4.**
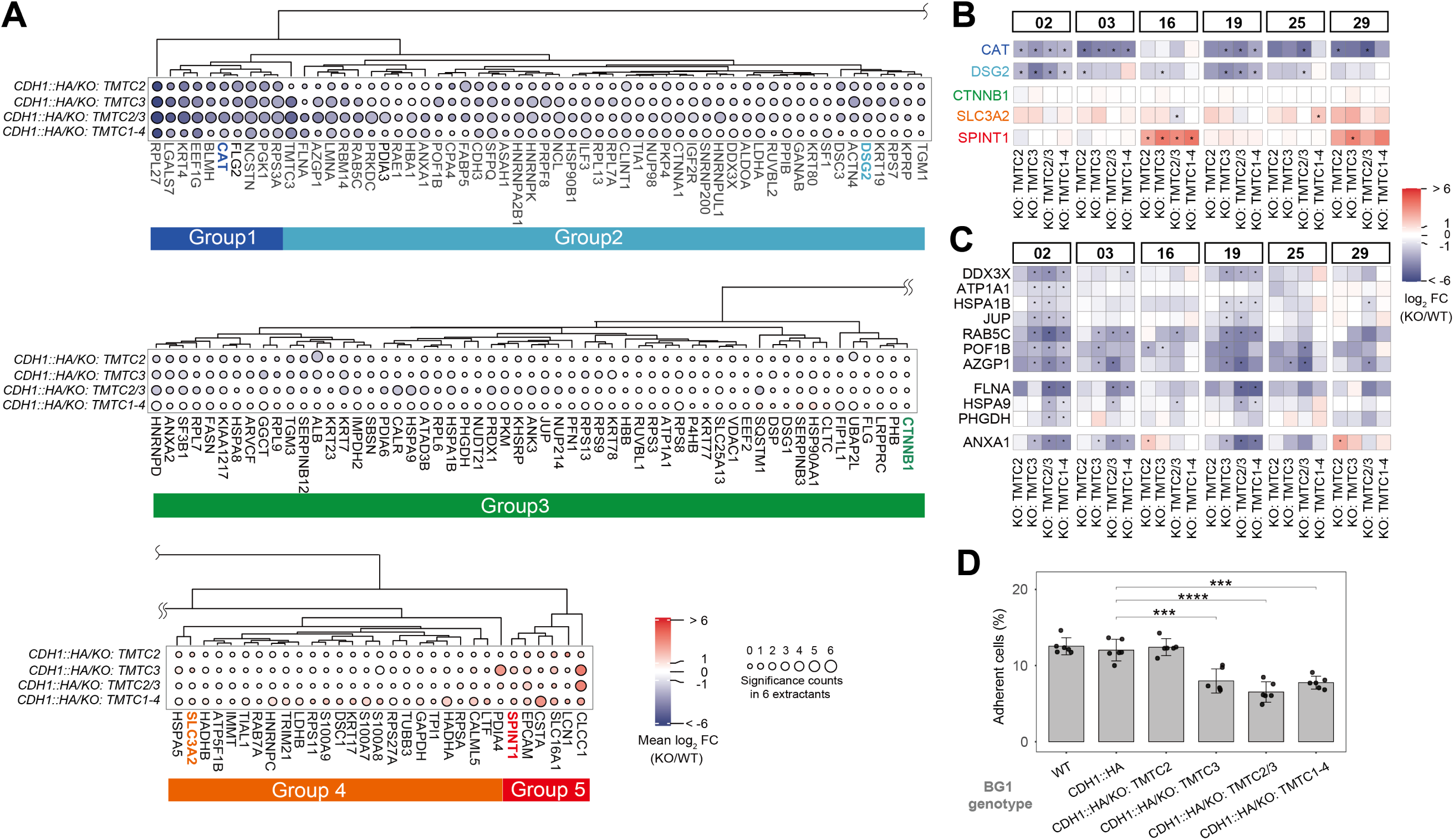
*O* -Man-dependent CDH1 interactome. **(A)** *Log_2_ fold change (FC) values of CDH1 interactors in TMTC deficient cell lines*: colors represent the average log_2_ FC values across the six IP conditions, with red indicating increased interaction with CDH1 and blue indicating decreased interaction with CDH1. Circles indicate the number of different IP conditions where the interactor was significantly changed in different *TMTC* KO cell lines (log_2_ FC ≥1 or ≤ −1, and p-adj. value ≤0.05). **(B)** Log_2_ FC values for selected interactors across different co-enrichment groups, for distinct IP conditions. Symbols indicate statistical significance, with ‘*’ representing log_2_ FC ≥1 or ≤ −1 and p-adj. value ≤0.05. **(C)** *Specific O-Man sites affect CDH1 interactions*: (upper-panel) example interactions affected by *O* -Man on EC domain G-strands; (middle-panel) example interactions affected by *O* -Man on EC domain B- and G-strands together; (lower-panel) ANXA1 exhibits increased co-enrichment when *O* -Man is depleted on EC domain B-strands (see conditions 16 and 20, KO: *TMTC2*). (D) Cell adhesion ability in cell lines expressing CDH1 with varying *O* -Man modification statuses; these data are presented as means ± SEM (n = 6). Statistical significance is denoted as follows: ***p ≤ 0.001; ****p ≤ 0.0001.

To interrogate the possibility that changes in enrichment of physical interactors within the CDH1 IPs can be explained by TMTC*-KO* -induced changes in the steady state abundances of putative interactors (rather than changes in biochemical affinity), we compared the proteomes of clarified cell extracts from each *TMTC* KO cell line to the *TMTC* WT. The abundances of most interactors remained unchanged when *TMTCs* were knocked out (Supp. Fig. 9); CDH3 was a notable exception, which showed ≳2-fold abundance decreases in *TMTC2*, *TMTC3*, and *TMTC1-4* KOs. GO analysis of the CDH1 protein interactions affected by *TMTC* KOs revealed distinct ontological enrichments (Supp. Fig. 10): interactors that exhibited decreased co-enrichment with CDH1 upon *TMTC* KO (Fig. 4A, Groups 1 and 2) were collectively enriched in annotations related to epithelial structure and function, including epidermis development, keratinocyte differentiation, and cell-cell junction organization. Additionally, these proteins were significantly associated with numerous RNA-related annotations. For the interactors exhibiting increased co-enrichment upon *TMTC* KO (Fig. 4A, Groups 4 and 5), antimicrobial immune-related annotations were prominently enriched, as were those connected with peptidase activity.

### Spatial *O* -Man glycans modulate CDH1 interactions and functions

Several proteins exhibit comparable changes in co-enrichment trends across the *TMTC* KO panel, which suggests that physical interactions with CDH1 may be more influenced by the presence or absence of *O* -Man and less by the specific modified EC domain B- or G-strand. For example, CAT (in Group 1) and DSG2 (in Group 2) exhibit this behavior (Fig. 4B & Supp. Fig. 8). In contrast, some interactions are apparently influenced by specific *O* -Man sites. Co-enrichment of DDX3X, ATP1A1, HSPA1B, JUP, RABC5, POF1B, and AZGP1 showed no change when *O* -Man glycans on the B-strands were depleted (*TMTC2* KO); yet these showed reduced co-enrichment with CDH1 when *O* -Man glycans on the G-strand were depleted (*TMTC3* KO). In contrast, the interaction between ANXA1 and CDH1 increased when *O* -Man glycans on the B-strand were specifically depleted (BG1*^CDH1::HA/KO: TMTC2^*). Furthermore, the co-enrichment of FLNA, HSPA9, PHGDH with CDH1 is diminished only when *O* -Man glycans are depleted from both the B- and G strands (Fig. 4C). These findings support the notion that CDH1 interactions are influenced not only by the presence (or absence) of *O* -Man but also by the specific location of these glycans.

Given the known role CDH1 in cellular adhesion and that its interactors were enriched for the biological process GO annotation “cell-cell junction,” we evaluated the influence of *O* -Man ablation on adhesion (Fig. 4D). We observed differences in adhesion ability among the various *TMTC* KO cell lines, despite the consistent expression level of CDH1 (Supp. Fig. 11). The results revealed that BG1*^CDH1::HA/KO: TMTC2^* did not lose its adhesion ability, whereas the adhesion ability significantly decreased when *TMTC3* was knocked out (BG1*^CDH1::HA/KO: TMTC3^*). We also observed a loss of adhesion ability in BG1*^CDH1::HA/KO: TMTC2/3^* and BG1*^CDH1::HA/KO: TMTC1-4^* cells, but there was no statistically significant difference compared to BG1*^CDH1::HA/KO: TMTC3^*. This is consistent with previous research reporting that TMTC3 influences cell adhesion ability (32). Given that different TMTCs target different *O* -Man sites of CDH1, the cell adhesion assay results suggest that *O* -Man deposition on the G-strand of EC domains (mediated by TMTC3) is necessary and sufficient for maintaining CDH1 adhesion functions.

Notably, we identified CDH3 as an interactor of CDH1 and observed decreased co-enrichment of the CDH1-CDH3 interaction in IPs from *TMTC* KO cells (Fig. 4A, Group 2), suggesting that these type I classical cadherins engage in interactions that at least partially depend on *O* -Man glycans. Yet, the reduction in co-enrichment could also be explained by a decrease in CDH3 steady state abundance, rather than by a reduction of biochemical affinity for CDH1. Indeed, we observed that CDH3 abundance decreased in response to *TMTC3* and *TMTC1-4* KOs (shown by western blotting in Fig. 5A; by mass spectrometry in Supp. Fig. 9): suggesting that *O* -Man glycans may stabilize CDH3 against turnover or otherwise regulate *CDH3* expression level. No difference in steady state CDH1 abundance was observed in bulk proteomic measurements by western blot (Supp. Fig. 11) or mass spectrometry (Supp. Fig. 9).

**Figure 5.**
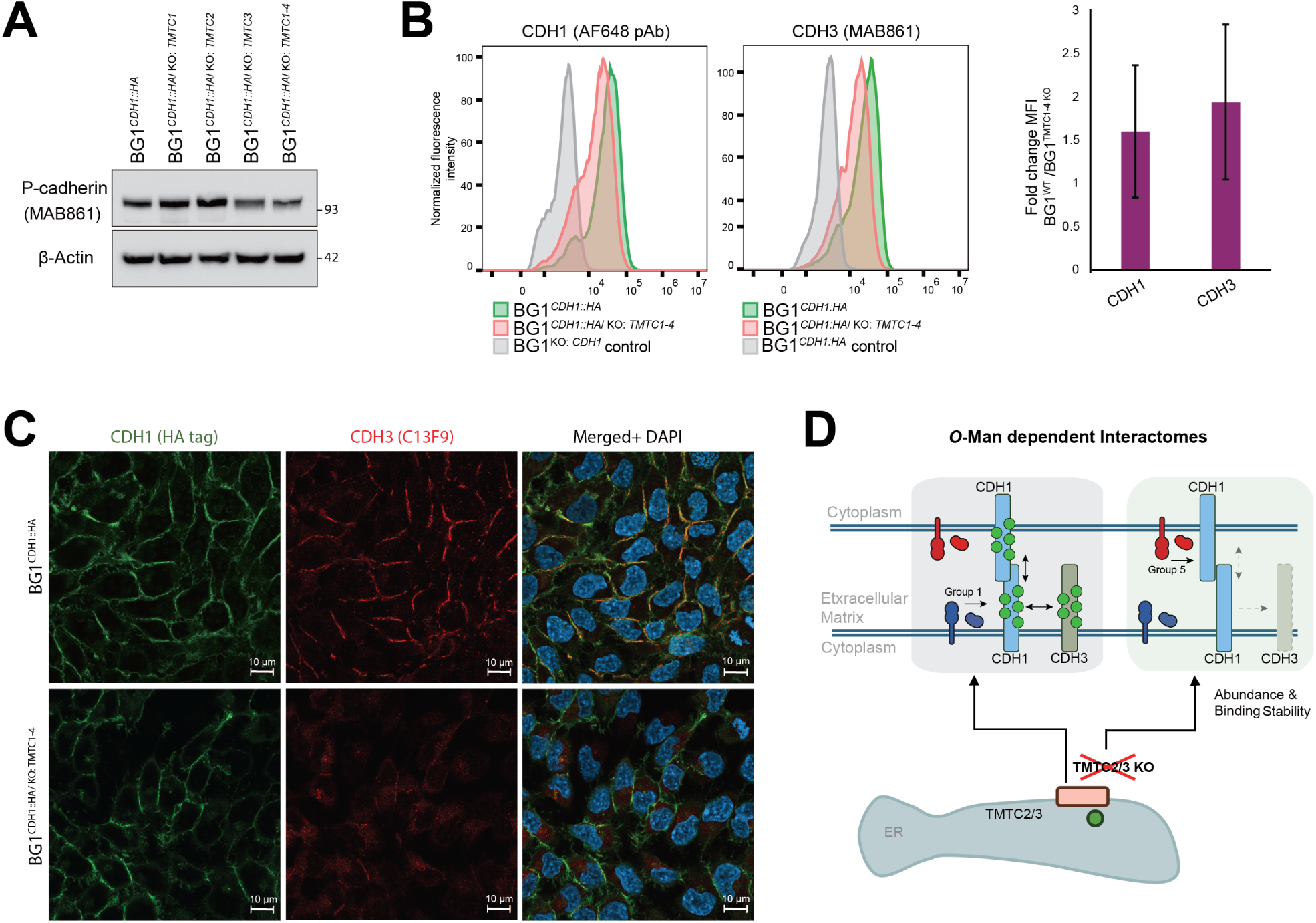
Effects of *TMTC* knockouts on CDH1 and CDH3 abundance and localization. **(A)** Western blot analysis of endogenous CDH3 abundance in BG1 cells with different *TMTC* KO statuses. **(B)** *Flow cytometry analysis of cell surface CDH1 and CDH3* : (left) representative histograms comparing fluorescence intensities in BG1*^CDH1::HA^* cells (green), BG1*^CDH1::HA/KO: TMTC1-4^* cells (pink), and BG1*^KO: CDH1^* negative control cells (grey); signals normalized to mode; (right) Quantification of fold-change in median fluorescence intensity for surface CDH1 and CDH3 in BG1*^CDH1::HA^* cells relative to BG1*^CDH1::HA/KO: TMTC1-4^* cells (n = 3). **(C)** Representative immunofluorescence images showing cellular localization of CDH1 (green) and CDH3 (red) in control BG1*^CDH1::HA^* cells (top panels) and BG1*^CDH1::HA/KO: TMTC1-4^* cells (bottom panels). Nuclei were counter-stained with DAPI (blue). Scale bar = 10 *µ*m. **(D)** *Schematic model of the O-Man-dependent CDH1 interactome*: some CDH1 interactors are modulated by *O* -Man, leading to decreased (Group 1) or increased (Group 5) co-enrichment based on changes e.g., in their affinity for CDH1, their localization and/or abundance (such as CDH3) upon *O* -Man depletion; *O* -Man depletion also reduces cell-cell adhesion.

To further interrogate putative CDH1-CDH3 interactions, their fractions at the cell surface were assayed by flow cytometry in BG1*^CDH1::HA^* and BG1*^CDH1::HA/KO:TMTC1-4^* cells. From this, we observed a ∼38% decrease in CDH1 and ∼50% decrease in CDH3, on average, in their respective cell surface abundances in *TMTC1-4* KO cells (Fig. 5B). Since the primary antibodies used for flow cytometry are specific for the EC domains of CHD1 and CDH3, we could not exclude the possibility that binding of these reagents may be influenced by *O* -Man. We therefore elected to visualize CDH1 by immunofluorescence using an anti-HA antibody (3F10) targeting its C-terminal HA-tag. Likewise, for CDH3, we used an antibody targeting its cytoplasmic domain (C13F9); the results are displayed in Figure 5C. We note, comparable results were obtained by immunofluorescence for CDH3 using both MAB861 and C13F9 (Supp. Fig. 12). In both BG1*^CDH1::HA^* and BG1*^CDH1::HA/KO:TMTC1-4^* cells, we observed a comparable, characteristic localization pattern of CDH1 at the interface of adherent cells, indicating that CDH1 cell surface localization is not significantly compromised by a lack of *O* -Man glycans. In contrast, while BG1*^CDH1::HA^* cells displayed overlapping signals for CDH1 and CDH3, BG1*^CDH1::HA/KO:TMTC1-4^* cells displayed reduced CDH3 signal at the cell surface, while apparent intracellular CDH3 was observed (Fig. 5C, Supp. Fig. 12). These results indicate that *O* -Man glycans may influence the local organization and stability of CDH3 molecules at the cell surface, rationalizing the observed decrease in the steady state abundance of CDH3 and reduced co-enrichment of CDH1-CDH3 interactions by IP, upon *TMTC* KO. Taken together, we summarize our findings using a schematic model, presented in Fig. 5D. This model proposes that *O* -Man on CDH1 is a key regulator of its protein interactions, modulating their steady state prevalence, to either promote (*e.g*., proteins in Group 1, Fig. 4A, are reduced on *O* -Man loss) or mitigate (*e.g.*, proteins in Group 5, Fig. 4A, are increased on *O* -Man loss) their accumulation (as quantified in this study by their co-enrichment in CDH1 IPs); this includes CDH1-CDH1 interactions (assayed by cell adhesion changes) and CDH1-CDH3 interactions (assayed by IP and IF), in which both interactors may be *O* -mannosylated.

## Discussion

In this research we have sought to identify the protein interactions associated with CDH1 and the degree to which those interactions are mediated by *O* -Man. For this, we employed endogenously HA-tagged CDH1 in the BG1 model cell line, glycoengineered with combinatorial KOs of *TMTC1-4* genes, combined with IP-MS based interactome screening (Fig. 1) and the I-DIRT signal detection method (Fig. 2). This provided us with (i) a focused list of steady state CDH1 interactors in BG1 cells (Figs. 2 & 3) and (ii) the opportunity to interrogate whether *O* - Man PTMs mediate those interactions, using IP-MS based interactome screening and label-free quantitative proteomics (Fig. 4A-C). We undertook differential *O* -Man glycoproteomic analysis to confirm complete ablation of *O* -Man on cadherins/protocadherins in *TMTC1-4* KO cells (Supp. Fig. 7) and we also assayed for loss of cell adhesion associated with *O* -Man ablation by *TMTC* KO (Fig. 4D). We observed that *TMTC3* KO was sufficient to compromise BG1 cell adhesion; this observation is consistent with the notion that CDH1-mediated cell adhesion may be regulated by *O* -Man without altering its absolute expression-level (Supp. Figs. 9 & 11). Noting that CDH3 was among the specific interactors exhibiting reduced co-enrichment with CDH1 upon *O* -Man loss (Fig. 4A), we further interrogated the molecular details of this change (Fig. 5).

### Logic and lessons from interactome screening

Extracting intact macromolecular assemblies associated with transmembrane proteins, like CDH1, is a well-known technical challenge (65, 66); they exhibit variable hydrophilicity across different domains, affecting solubility upon extraction, and the maintenance of interactions with phospholipids may be critical to the maintenance of native folds and protein interactions. Drawing from previous experience (46, 47, 57, 67), we systematically evaluated the effects of different reagents on (i) the yields of affinity enriched CDH1 and (ii) the profiles of co-enriched proteins (i.e., putative interactors). We observed that, at typical working concentrations, the detergents Triton X-100, CHAPS, DDM, and C12E8 aided CDH1 affinity enrichment, whereas Tween 20 and Brij 58 did not (Fig. 1C & Supp. Table 3). We also observed that extraction solutions containing potassium acetate (including extractants 2, 3, 15, 18, 19, 22, 23, etc., in Fig. 1C & Supp. Table 4) co-enriched more proteins; however, this salt also appeared to increase presumed non-specific contaminants (based on our visual inspection criteria and corroborated by I-DIRT analysis: compare extractants 2, 3, and 19 to the others in Fig. 2B).

When moving from the pre-screening stage to a reduced set of screening conditions, the liberated bandwidth is used to include multiple replicates for each condition explored, enabling more sophisticated quantitative analyses. For the condition selection process, we prioritized features that are readily observed in stained gels, e.g., lanes that exhibited: (i) relatively discrete patterns of sharply stained bands (i.e., potential components of a stable assembly), (ii) a relative paucity of faint/fuzzy background staining (i.e., potential post-extraction noise, especially when present across a very broad mass range), and (iii) a high intensity band that corresponds to the target protein molecular mass. We also prioritize (iv) IP conditions exhibiting or missing, one or more distinctive, discrete bands; these may reveal dependencies on certain test tube solutions, for the maintenance of one or more interactions together with the target. We may select complex profiles with faint/fuzzy background staining, provided the target protein is highly abundant and some other bands in the lane are discrete and relatively intense. Among shortlisted screening conditions that moved to subsequent rounds, the above considerations were counterbalanced against maintaining high proteomic complexity (relative to all the conditions explored, to mitigate false negatives; i.e., considering all the information displayed in Fig. 1C & Supp. Fig. 2). Together, these criteria constitute an integration of quality assurance, discovery optimization, and the prudent use of downstream experimental bandwidth, including MS instrument time. We envision an objective, computer-automated selection process in future iterations.

To mitigate the inclusion of false positives (i.e., post-extraction artifacts) in our analyses, we turned to I-DIRT, which revealed that over half of the co-enriched proteins were potentially spurious interactors. Yet, some of these proteins may be false negatives - real interactors that rapidly decay and re-equilibrate in a suboptimal extraction solution. The use of multiple different extraction solutions, combined with I-DIRT, therefore enabled us to screen more comprehensively for specific CDH1 interactors, including both stable and labile/transient interactors (post-extraction), within the biochemical breadth of experimental conditions explored. Indeed, of the 27 specific interactors that we found in common with the prior proximity labeling-based studies (Fig. 2C), only 7 were identified as specific in ≥ 5 interactome screening conditions. 5 of the proteins in common between studies were identified as specific in only 1 (of 6 tested) interactome screening condition. These observations support our assertion that many IP conditions must be explored to identify those suitable for the post-extraction maintenance of native interactions that normally occur *in situ* (i.e., within myriad biochemical milieux present inside of cells), for any given target macromolecular assembly; likewise, broad exploratory screening also enables comprehensive rejection of post-lysis noise that may idiosyncratically accumulate on different macromolecular assemblies during their transfer to artificial solutions in test tubes. We note that certain common contaminants, such as keratins, were enriched in the I-DIRT label-swapped experiment (light-SILAC BG1*^CDH1::HA^*), while the corresponding ratio in I-DIRT (heavy-SILAC BG1*^CDH1::HA^*) was near zero (Supp. Fig 6). This is explained by the fact that common exogenous contaminants originating from experimenter and environment contain only naturally occurring isotope compositions in their amino acids. However, some keratins, such as KRT4 and KRT17, were identified as specific interactors based on their I-DIRT ratios. Notably, of the TMTC isoenzymes, only TMTC3 was identified as a specific interactor in four of the I-DIRT experiments (Extractant 2, 3, 19, and 29); considering the usual difficulty to isolate enzyme:target interactions, we rationalize this result as follows: CDH1 is not expected to be a substrate of TMTCs 1 and 4; TMTC3 is the putative enzyme primarily responsible for CDH1 *O* -Man. TMTC3 adds multiple *O* -Man residues on G-strands of EC domains 2, 3, and 4 (up to ten *O* -Man sites); in contrast, TMTC2 adds one or two *O* -Man residues on the B-strands of EC domains 2 and 4 (13), potentially explaining a reduced avidity of this enzyme:target complex. Additionally, while TMTC3 was detected in our proteomic analyses of clarified BG1 cell extracts, TMTC1, 2, and 4 were not - indicating a lower general abundance of these proteins in BG1 cells (Supp. Fig. 11). Our ongoing studies (manuscript in preparation) show that CDH1 is targeted by TMTC2 and 3 (therefore *TMTC2/3* KO ablates all CDH1 *O* -Man); but other cadherins are also targeted by these enzymes and our mass spectrometry-based survey revealed that 10 different cadherins and protocadherins are affected by complete *TMTC* ablation in BG1 cells harboring *TMTC1-4* KO (assayed by mass spectrometry in Supp. Figure 7), in agreement with our recent study (55).

### Summary of CDH1 interactomics: limitations, findings, and future directions

Screening endogenously HA-tagged CDH1 allowed us to identify 148 members of a CDH1 interactome in BG1 cells (Fig. 2B); among these, we confirm 27 putative interactors, asserted by two earlier studies (Fig. 2 C, Supp. Table 5) (39, 40). IP-MS analyses using endogenous tagging avoids artifacts associated with studies using ectopic expression of tagged proteins: often, ectopic constructs are over-expressed from non-native promoters, exaggerating the abundance of the tagged protein-of-interest and thereby altering the constellation of *in vivo*binding partners readily assayed by MS; or, when the abundance of the tagged ectopic protein-of-interest is well calibrated to that of the endogenous protein, only a maximum of ∼½ of total pool is subject to capture and analysis (potentially limiting absolute sensitivity) (57). While the endogenously HA-tagged CDH1 cells used here avoid these limitations, we acknowledge that the impact of *O* -Man on *cis*- and *trans*-binding of CDH1-CDH1 homodimers and oligomers remains difficult to assess: in this context, all CDH1 molecules harbor the tag and are therefore potentially subject to capture by the *α*-HA antibody regardless of their glycosylation and interaction state; as such we could not quantify the alterations in CDH1-CDH1 interactions except by the surrogate cell adhesion assay (Fig. 4D; to the degree that cell adhesion relies on CDH1-CDH1 interactions). We envision that future experiments, e.g., involving endogenously tagged CDH1-HA and -FLAG, in different CRISPR-edited cells, could address *trans* alterations; whereas the same, in a single heterozygous CDH1^HA/FLAG^ cell line, could address *cis* alterations.

Among our results, we found that CDH1 can interact with other adhesion molecules (such as EPCAM, CDH3, DSG1/2, DSC1/3) and associated junctional proteins (like ARVCF, DSP, JUP); these maintain cell-cell junctions, are involved in epidermal cell differentiation, and facilitate intercellular communication (Fig. 3). Analysis of the *O* -Man-dependent CDH1 interactome revealed that many such interactions are apparently modulated by *O* -Man; the degree and direction of modulation (increase or decrease) varied (Fig. 4A-C). For some CDH1 interactors, such as CTNNB1, no effect of *O* -Man ablation was observed; this is rationalizable by the fact that *O* -Man decorates EC-domains (presented in the extracellular matrix), whereas CTNNB1 binds to the C-terminus of CDH1 (which is presented to the cytosol) (23). Yet, many interactors were affected by *O* -Man ablation, exhibiting increased or decreased co-enrichment with CDH1.

For example, our results identify EEF1G, BLMH, CAT, and FLG2 as TMTC1-4 dependent cytoplasmic interactors of CDH1, and while we still lack molecular insight and explanations for these observations, it is interesting to note that cytoplasmic proteins categorized as Group 1 CDH1 interactors (Fig. 4A) exhibit this behavior, indicating that extracellular PTMs influence interactions with cytoplasmic partners. FLG2, encoding the 248 kDa cytoplasmic protein filaggrin-2, has been suggested to promote cell-cell adhesion in the upper epidermis through unknown mechanisms implicating classical and desmosomal cadherins (68, 69), which is supported by our observations of TMTC1-4 dependent CDH1-FLG2 interaction. Further studies, guided by our interactome screen, are clearly needed to validate and systematically address the molecular connection between CHD1 and such cytoplasmic partners. In contrast, the apparent stoichiometry of other interactors increased upon depletion of O-Man (Group 5 in Fig. 4A). This group includes SPINT1, EPCAM, and CSTA, which are primarily involved in maintaining epithelial integrity, regulating cell adhesion and signaling (EPCAM) (70–72). The enhanced association with these functionally diverse proteins suggests that *O* -Man is a key regulator of cellular and tissue homeostasis through multiple pathways.

Furthermore, the *O* -Man dependent CDH1 interactome showed that these interactions can be spatially specific regarding *O* -Man sites (Fig. 4C). Co-enrichment of ANXA1 with CDH1, for example, decreased upon loss of *O* -Man glycans on the G-strand but increased when *O* -Man was lost from the B-strand. ANXA1, a phospholipid-binding protein, is involved in diverse cellular processes. The disruption of the normal balance between ANXA1 and CDH1 function is strongly implicated in cancer progression; in many aggressive cancers elevated ANXA1 expression correlates with reduced CDH1 expression or function, induction of EMT, increased invasiveness, metastasis, and poor patient prognosis (73, 74) – revealing the likely importance of considering O-Man in addition to expression levels and mutations when investigating CDH1 function.

The macromolecular assembly of CDH1 interactors dynamically regulates CDH1 adhesive functions (75, 76). Cells are capable of inducing passive and active adhesion states for CDH1 at the cell surface through *inside-out* signaling events, akin to integrin activation (77–79), where binding of cytoplasmic partners induce conformational changes that are transmitted to extracellular CDH1 EC-domains which influence CDH1 adhesive functions on the opposite side of the plasma membrane (75, 76). Such passive and active CDH1 states can be induced by “activating” antibodies that bind distinct extracellular CDH1 epitopes, thus stabilizing conformations favoring the active adhesive state of CDH1, which concomitantly triggers a biochemical event in the opposite, outside-in direction, resulting in dephosphorylation catenin delta-1 (CTNND1), a key binding partner at the cytosolic domain of CDH1 (30, 79–81). Our results, identifying new *O* -Man dependent CDH1 interactors, further support this model and indicate that CDH1 EC-domains may require *O* -Man at distinct sites to fine-tune allosteric regulation, conformational states, and interactions of CDH1. We reason that EC-domain O-Man glycans are not necessarily (or exclusively) involved in direct protein-glycan interactions but may act as structural elements that are required to balance EC-domain stability and flexibility, and thereby CDH1 macromolecular assemblies and functions.

The precise molecular mechanism underlying these observations are beyond the scope of this study; what we believe is apparent from this work, however, is that specific cadherins have *O* -Man dependent functions, which fine-tune cell surface organization, and by extension, cell surface interactomes. Future research directions could focus on a deeper functional analysis of glycosylation-dependent interactions and their possible roles in disease onset and/or progression. The identified interactors and their modification-dependent binding patterns may represent therapeutic opportunities or biomarkers for diseases associated with CDH1 dysfunction. We believe the optimized protocols and analytical strategies developed herein constitute a robust framework for investigating integral membrane protein interactomes and their PTM-dependencies. Given this proof-of-principle success with CDH1, we anticipate expanding our landscape of such targets using these methods and hope the field will do the same.

## Acknowledgements

We thank Francis Jacob (University Hospital Basel) for providing BG1 cells. We thank Karin Wolters (UMCG), Ida Signe Bohse Larsen and Lorenzo Povolo (University of Copenhagen), for experimental support and constructive discussions; and we thank Lianli Chi (Shandong University) for his generous support of SX’s China Scholarship Council (CSC) project.

## Author Contributions

Conceptualization: JL, AH; Data Curation: SX, KM; Formal analysis: SX, OGRB; KM, SYV; Funding acquisition: JL, AH, SX; Investigation: SX, KM, WT; Methodology: JL, AH, PH, SX; Project administration: JL, AH; Resources: JL, AH, PH; Software: SX, OGRB; Supervision: JL, AH, PH, HP; Validation: SX, KM; Visualization: SX, JL, KM, AH; Writing—Original Draft: SX, JL, KM, AH, WT; Writing—Review & Editing: SX, JL, KM, AH, PH

## Data availability

Mass spectrometry-based proteomics data produced in this work have been deposited in the MassIVE database with identifier MSV000097524. The in-house developed R-code is available at https://github.com/XieShaoshuai/E-Cadherin-interactome.

## Supplemental Figures

**Supplemental Figure 1.**
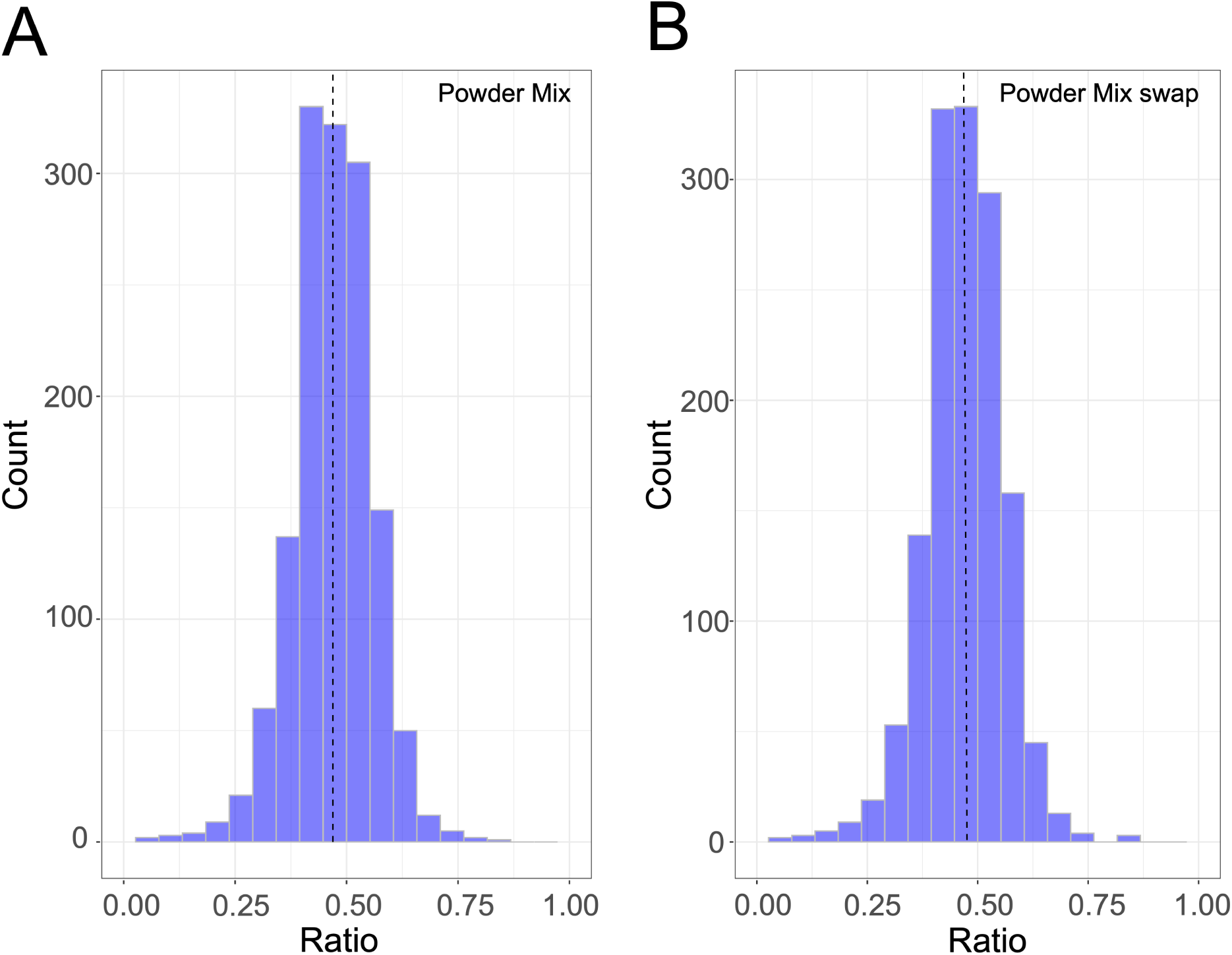
Ratio Distribution of Cell Powder Mix. **(A)** Ratio distribution for cell powder mix (light-labeled BG1^WT^ and heavy-labeled BG1*^CDH1::HA^*). The median ratio for all proteins across 6 replicates is 0.45. The standard deviation (SD) of the median ratio among six replicates is 0.0017. **(B)** Ratio distribution for swapped cell powder mix (heavy-labeled BG1^WT^ and light-labeled BG1*^CDH1::HA^*). The median ratio for all proteins across 6 replicates is 0.48. The standard deviation (SD) of the median ratio among six replicates is 0.0012.

**Supplemental Figure 2.**
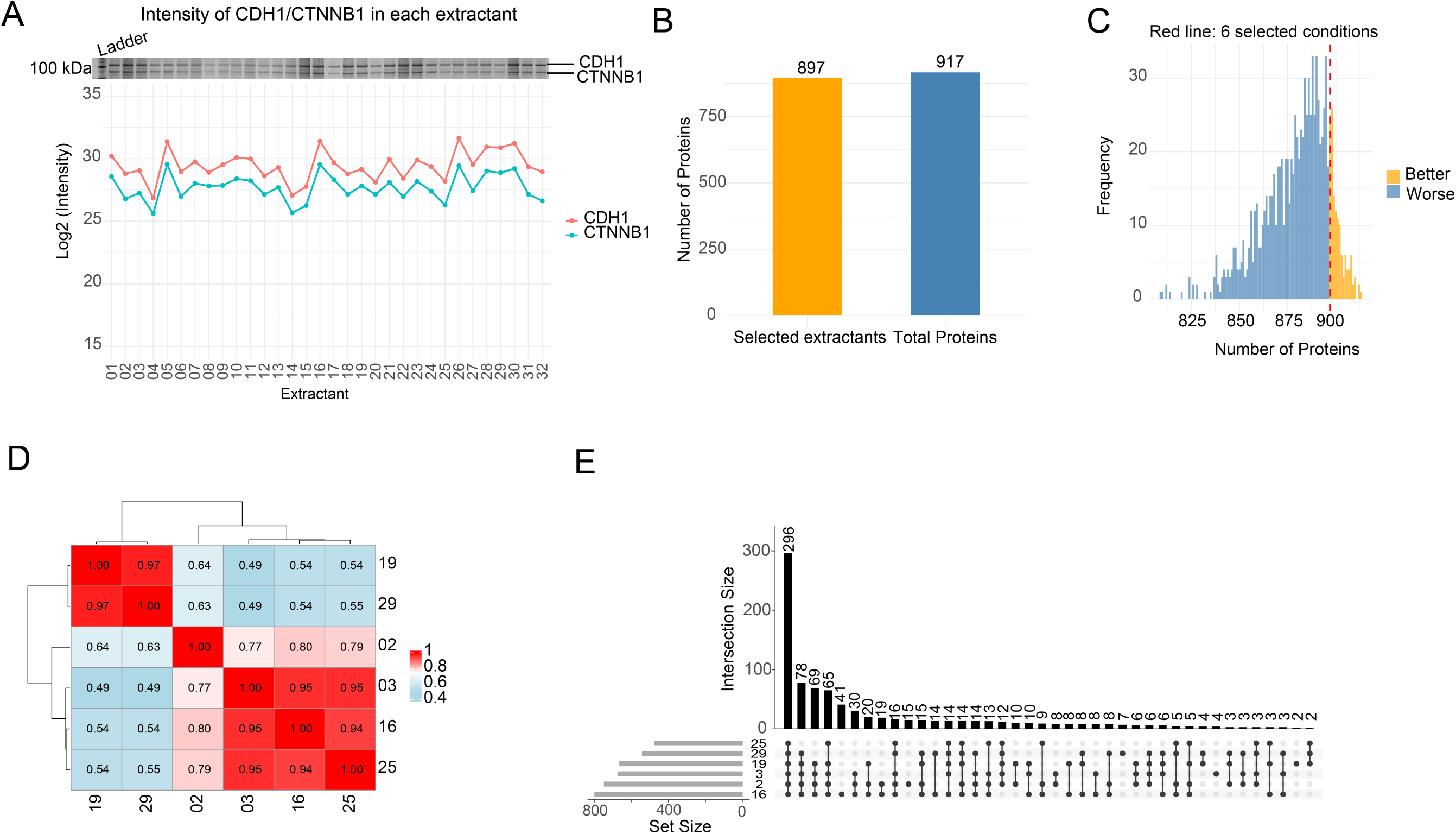
Extractant Selection for IP Screening. **(A)** CDH1 and its interactor CTNNB1 intensities in the SDS-PAGE gel (silver stain) and label-free mass spectrometry. **(B)** Number of proteins identified for selected 6 extractants and all 32 extractants. A total of 917 proteins were identified in 32 extractants, while 897 proteins were identified in the 6 selected extractants. **(C)** Distribution of identified protein numbers from 1000 random selections of 6 extractants. The number of proteins for the 6 selected extractants in Fig. 1C is indicated by a red dotted line. **(D)** Correlation of LFQ MS results for 6 selected extractants. **(E)** UpSet analysis of proteins identified in the 6 selected extractants. All ms-based analyses use protein intensity values output from Proteome Discoverer (“Abundance”), as per Fig. 1C.

**Supplemental Figure 3.**
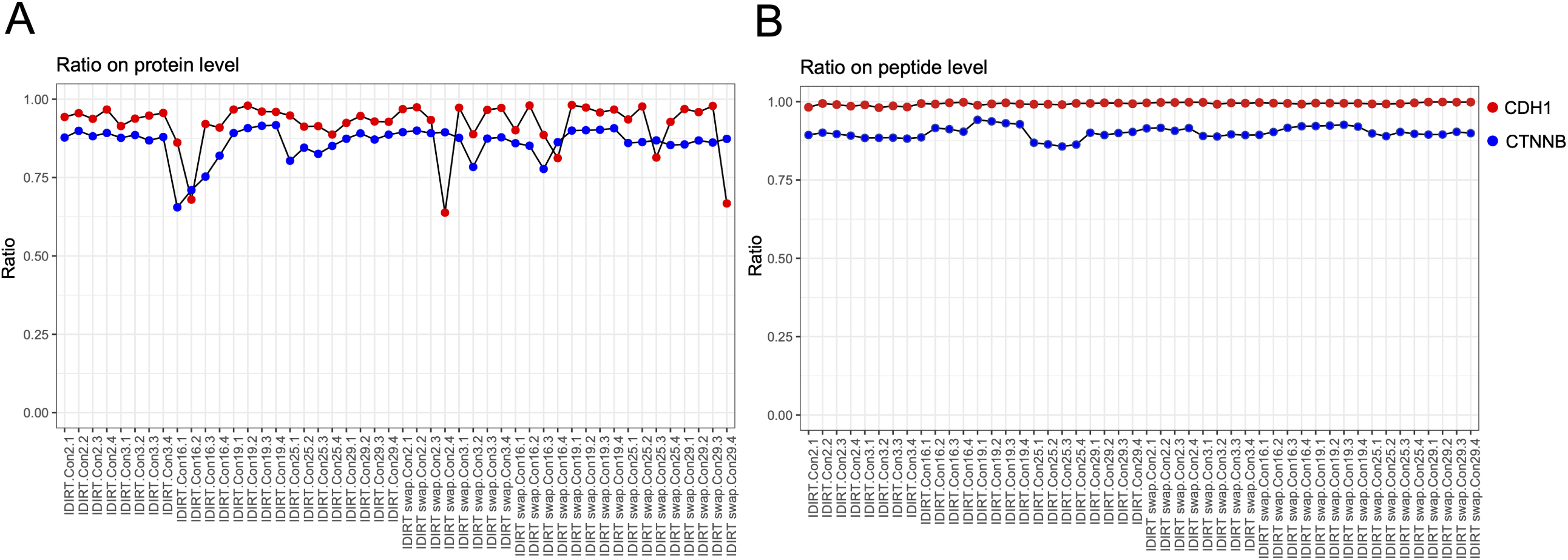
I-DIRT Ratios for CDH1 and CTNNB1. **(A)** I-DIRT ratios for CDH1 and CTNNB1 based on protein intensity, quantified using Proteome Discoverer. **(B)** I-DIRT ratios for CDH1 and CTNNB1 based on peptide intensities, quantified using Proteome Discoverer.

**Supplemental Figure 4.**
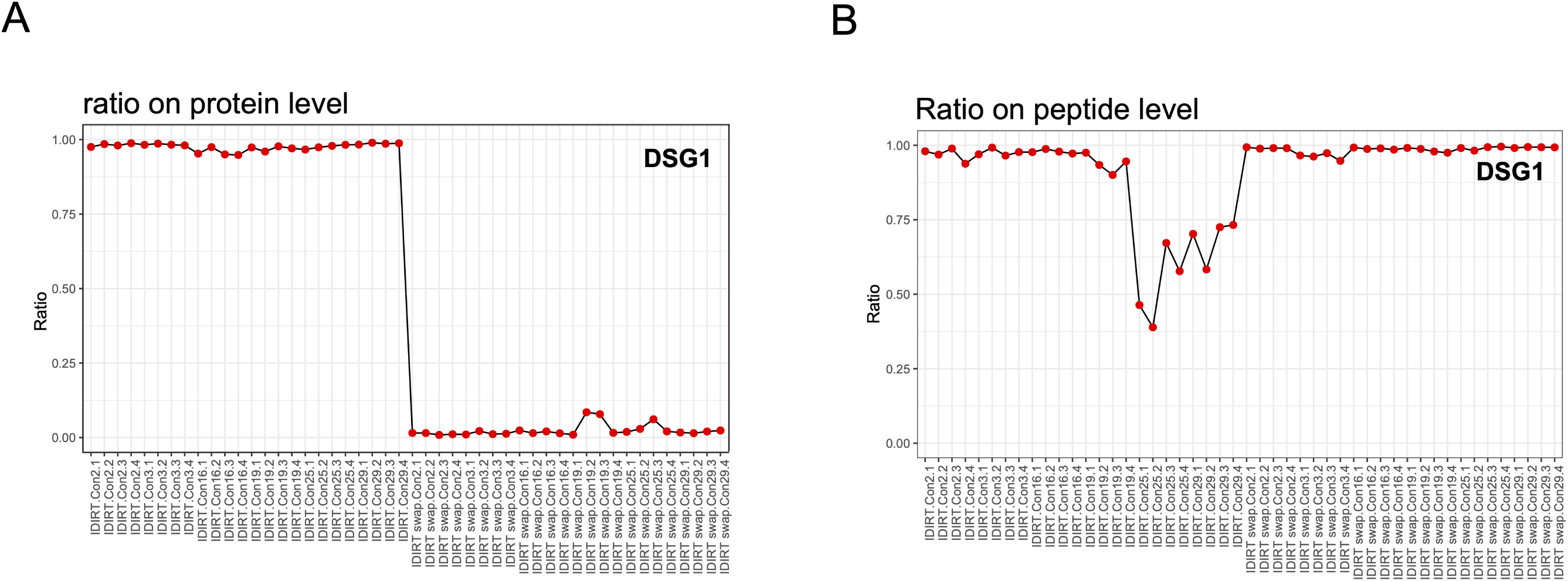
I-DIRT ratio of DSG1. **(A)** I-DIRT ratios for DSG1 based on protein abundance quantified by Proteome Discoverer. **(B)** I-DIRT ratios for DSG1 based on peptide abundance quantified by Proteome Discoverer.

**Supplemental Figure 5.**
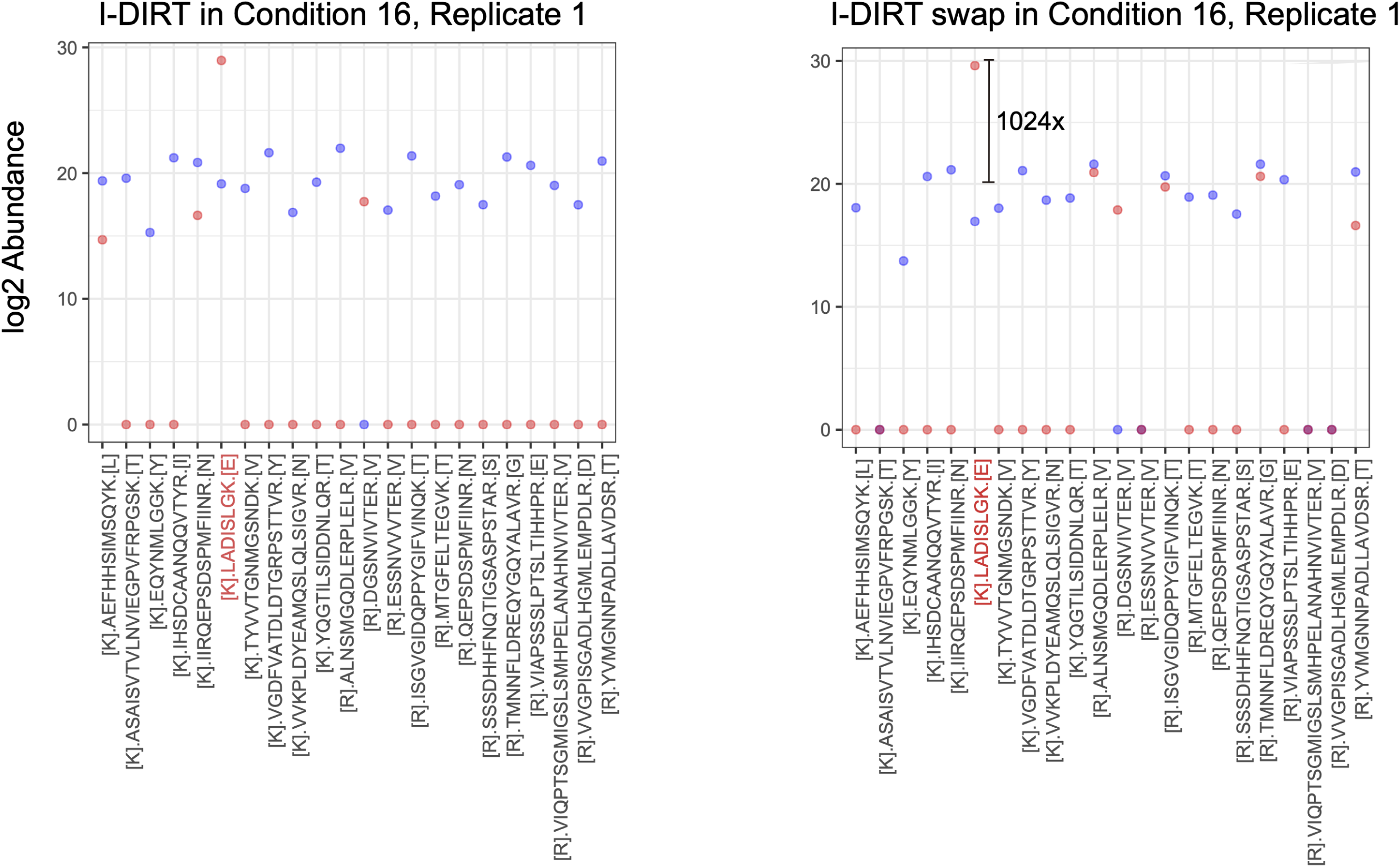
Peptide abundance of DSG1 in I-DIRT experiments. One replicate each for condition 16 in I-DIRT and I-DIRT swap experiments is shown respectively. The abundance of a single heavy-labeled peptide (red) is nearly 1000 times higher than that of other peptides, disproportionately contributing to the total protein abundance.

**Supplemental Figure 6.**
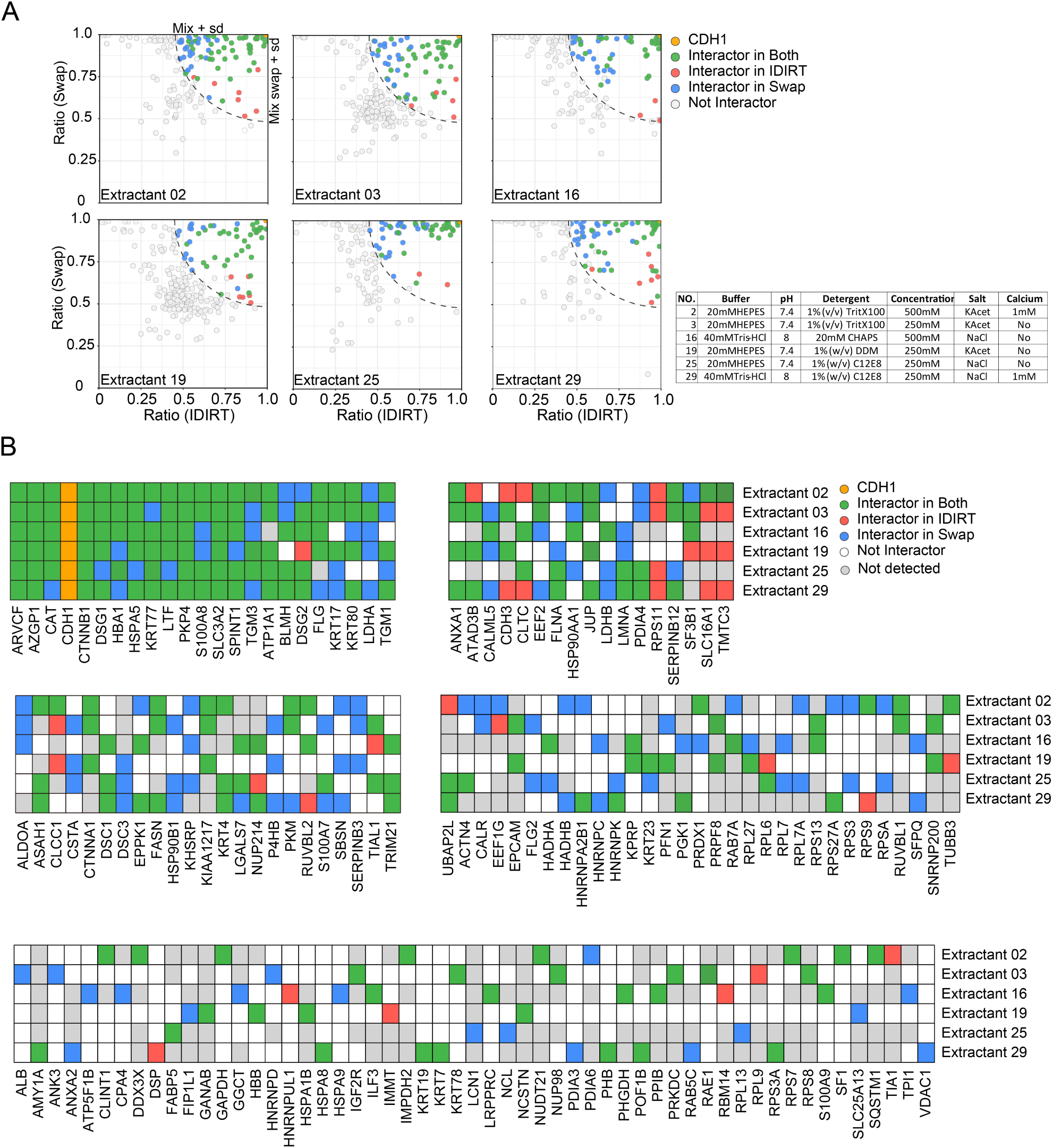
I-DIRT specific CDH1 interactors identified in different IP conditions. **(A)**Specific interactors were identified by comparing protein enrichment ratios from the I-DIRT and I-DIRT swap experiments. For each experiment, a Bayesian test was applied to compare the enrichment ratio of a given protein to the median ratio of all proteins in the corresponding bulk proteome (mix or mix swap). Proteins were classified as specific interactors if they met two criteria: a Bayesian Factor (BF) > 3 and location within the significance ellipse. The ellipse’s semi-major and semi-minor axes were defined by the median ratio plus or minus one standard deviation (SD) of the bulk proteome, respectively. The table on the right shows the formulation of the selected six extractants. **(B)** Summary of the CDH1 interactome across six IP extractants.

**Supplemental Figure 7.**
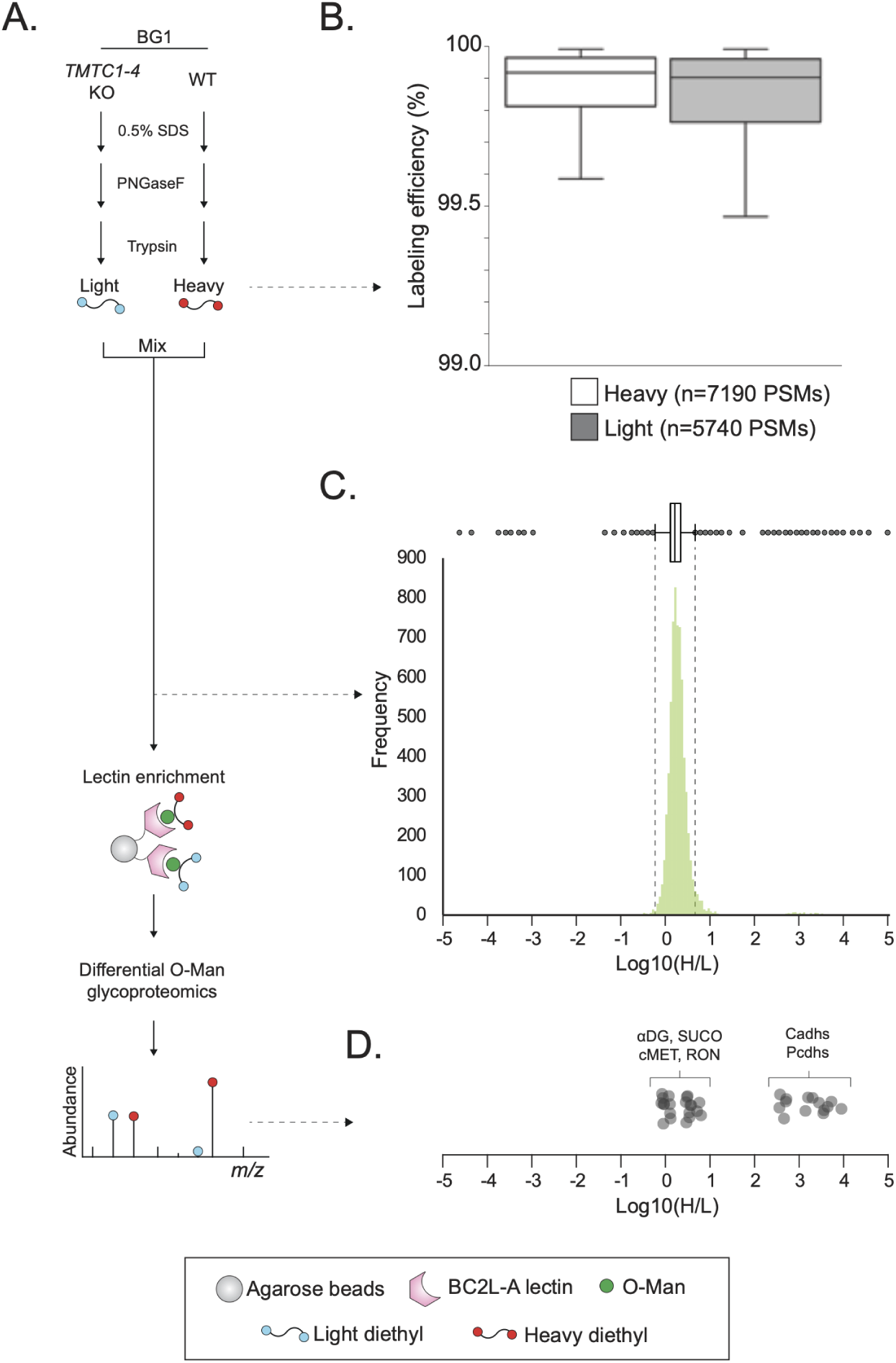
*TMTC1-4* dependent *O* -Man glycosylation of cadherins and protocadherins in BG1 cells. **(A)** Workflow for comparative analysis of O-Man glycosylation in BG1^WT^ and BG1*^KO: TMTC1-4^* cells, as previously described (55). **(B)** Diethyl stable isotope (DEL) labeled total cell tryptic digests were individually analyzed and MS1 peaks of peptide spectral matches (PSMs) were used to assess labeling efficiency (EL), calculated by EL = 1-(1/((labeled/unlabeled) +1))). **(C)** Heavy (H) and light (L) labeled tryptic digest were mixed (volume 1:1) and analyzed to assess mixing ratio and proteome variability. The bar chart shows log10 transformed H/L frequencies of identified peptides (n=6664 PSMs). Outliers were determined by Q1-1.5xIQR and Q3+1.5xIQR boundaries (dashed lines). **(D)** BC2L-A lectin enriched fractions were analyzed as previously described (55), and relative abundances of identified *O* -Man glycopeptides plotted as log10(H/L) ratios. The dot-plot shows a group of glycopeptides (<10-fold change) originating from αDG and SUCO (POMT1/POMT2 substrates) as well as cMET and RON receptors (TMEM260 substrates), while the canonical TMTC1-4 substrates including cadherins (Cadhs) and protocadherins (Pcdhs) show >100-fold change, thus demonstrating that *TMTC1-4* KO abolishes cadherin-specific *O* -Man glycosylation in BG1 cells.

**Supplemental Figure 8.**
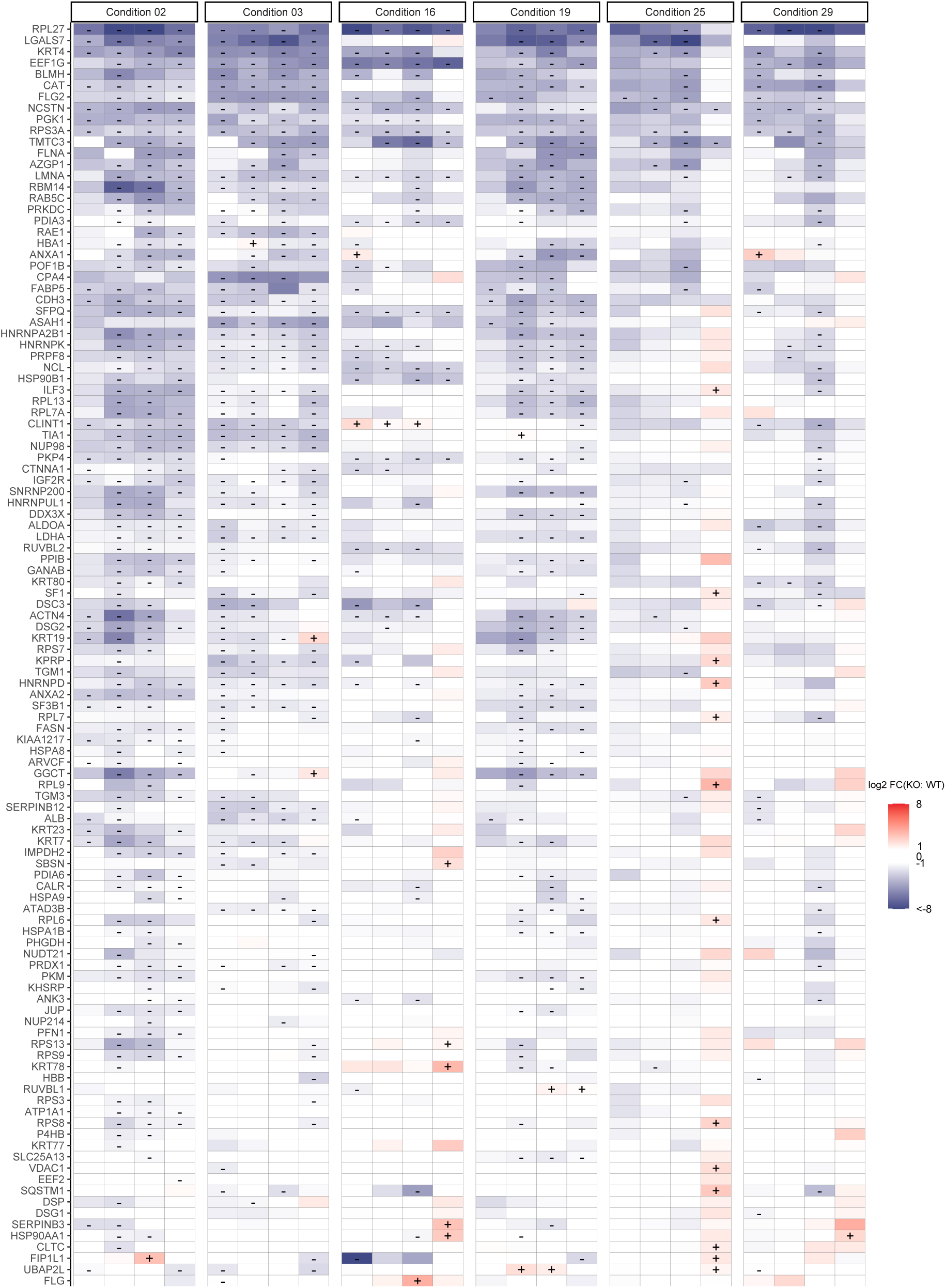
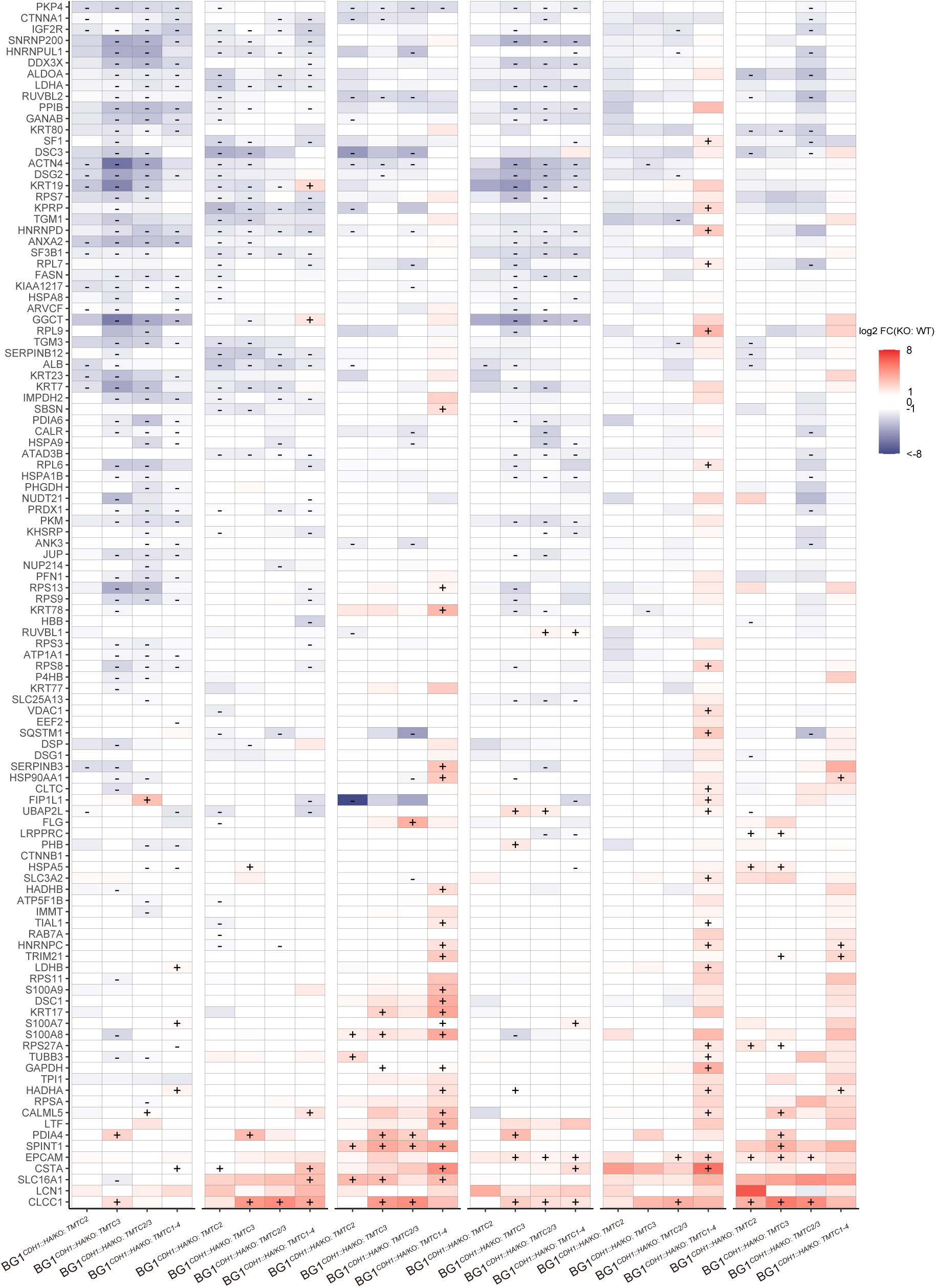
The *O* -Man dependent E-cadherin interactome. Heatmap of log_2_FC of CDH1 interactors across four *TMTC* KO cell lines under six IP conditions. Rows display interactors, while display IP conditions for each knockout cell line. Red indicates increased enrichment and blue indicates decreased enrichment, with color intensity reflecting the effect size. Symbols indicate statistical significance, with “+/−” representing log_2_FC >1 or < −1 and p-adj. ≤ 0.05.

**Supplemental Figure 9.**
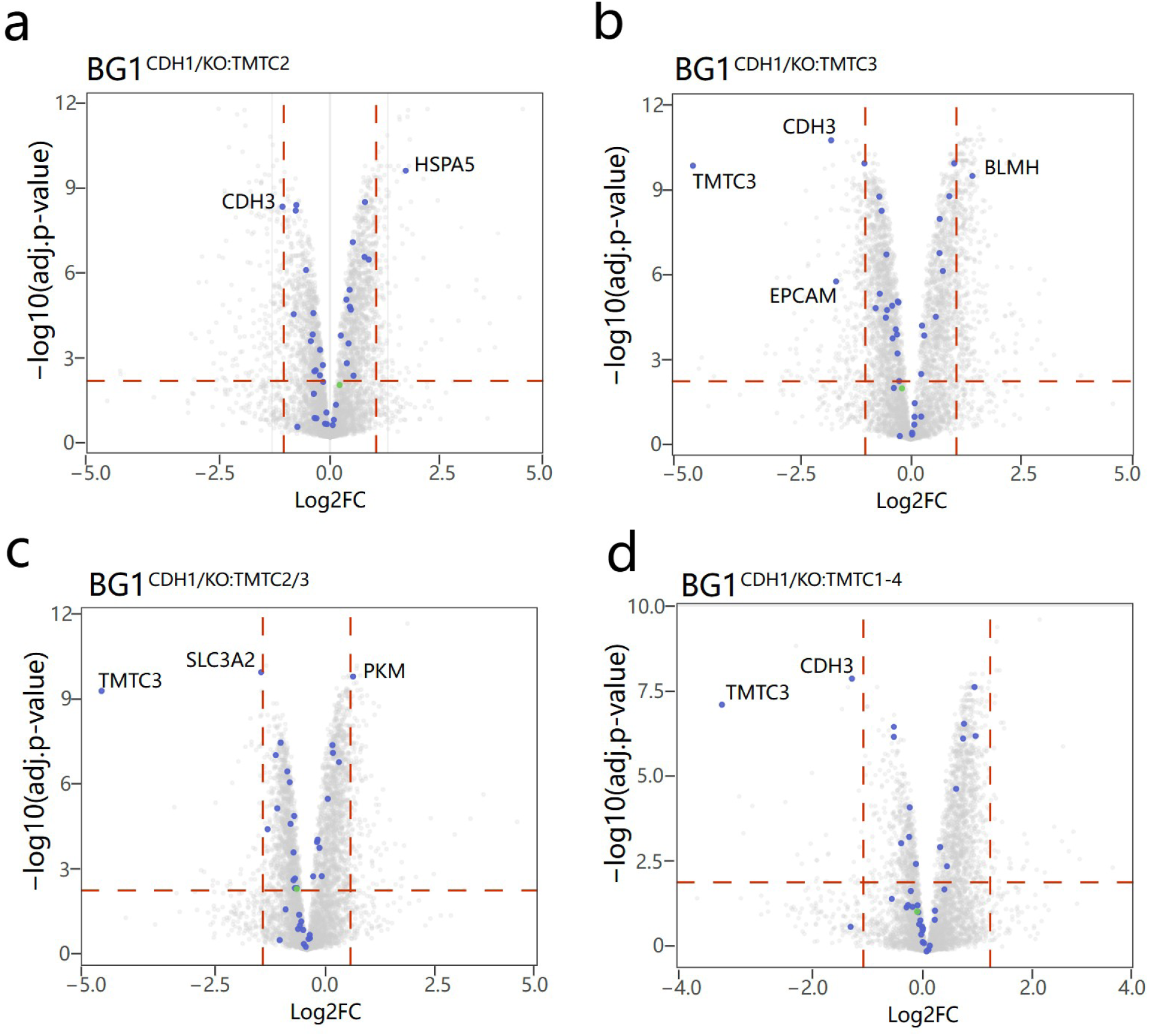
Whole cell lysate proteome analysis for different cell lines. The I-DIRT specific CDH1 interactors are colored blue. CDH1 is colored green. Only proteins with abundance changes in the given cell extracts are labeled with the Uniprot gene symbol.

**Supplemental Figure 10.**
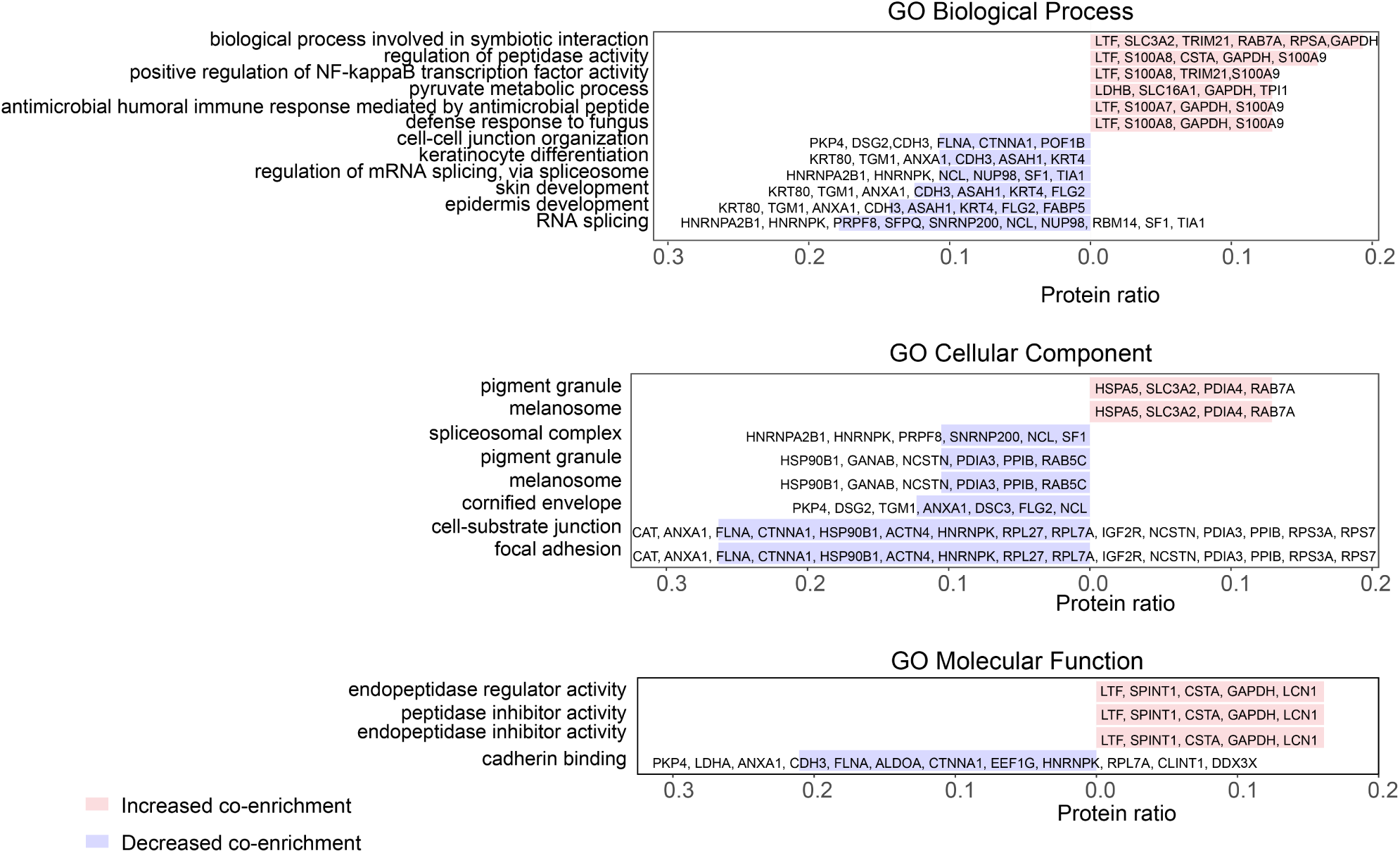
Gene Ontology (GO) analysis was performed on CDH1 interactors identified as O-Man-dependent. The results show enriched terms for Biological Process (BP), Cellular Component (CC), and Molecular Function (MF), highlighting the key roles of these interactors in adhesion, signaling, and protein stability. Categories are presented with a protein ratio greater than 10% and a p-value < 0.01.

**Supplemental Figure 11.**
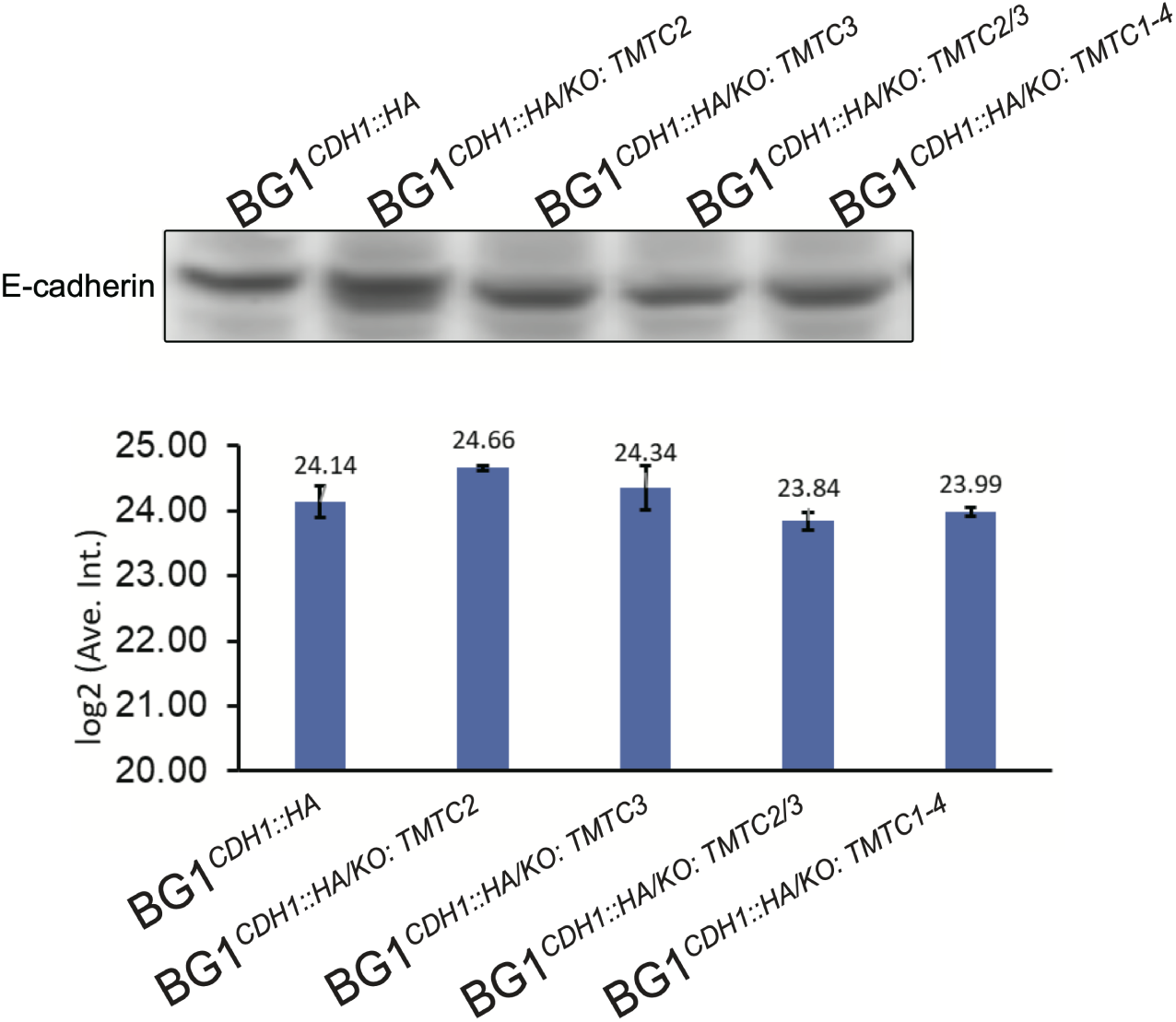
Expression of E-cadherin in different cell lines. The expression of E-cadherin was quantified via Western blot analysis (n = 3). 20 *µ*g of lystates were loaded in each lane. 20 *µ*g of cell lysate (processed following *Bulk protein extraction from cells*, see *Methods*) was diluted in 1× LDS (Invitrogen), reduced with dithiothreitol (10 min, 90*^◦^*C), and resolved on NuPAGE Bis-Tris 4–12% gels (Invitrogen) using 1 × MOPS buffer (Invitrogen). Proteins were transferred (1.5 h, 90 V) to a methanol-activated PVDF membrane with Tris-Glycine transfer buffer (25 mM Tris, 192 mM glycine, 10% ethanol). The membrane was blocked for 1 h at room temperature (RT) with gentle agitation in TBST containing 5% (w/v) non-fat milk, followed by incubation with an HA-Tag Monoclonal Antibody (#26183, Thermo Fisher, 0.2 *µ*g/mL in TBST containing 5% (w/v) BSA and 0.02% (w/v) sodium azide). After washing with TBST, HA-tagged CDH1 was detected using an HRP-conjugated anti-mouse secondary antibody (Thermo Fisher, 1:10,000 dilution in TBST).

**Supplemental Figure 12.**
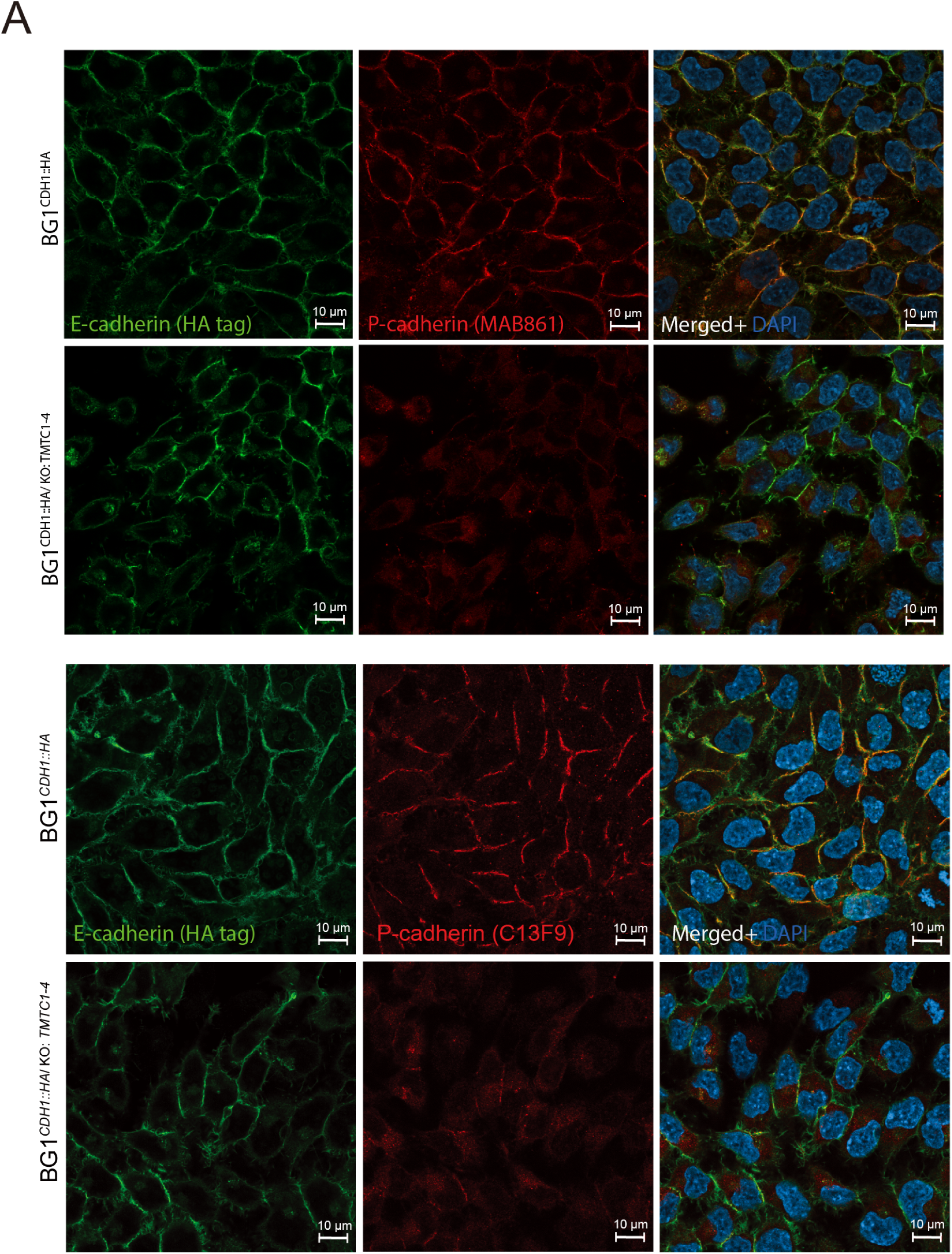
Comparison of CDH1 and CDH3 localization in WT and O-Man depletion cells. Cells were stained for CDH1 via its HA-tag (green; anti-HA antibody 3F10) and for CDH3 (red; anti-CDH3 antibody MAB861 or C13F9). Nuclei were counterstained with DAPI (blue). Scale bar = 10 *µ*m.

